# Synergistic and redundant information dynamics are modulated by Alzheimer’s disease and cognitive impairment

**DOI:** 10.64898/2026.02.18.706630

**Authors:** Keenan J. A. Down, Jonathan Huntley, Pedro A. M. Mediano, Daniel Bor, the Alzheimer’s Disease Neuroimaging Initiative

## Abstract

**Background:** The early diagnosis of Alzheimer’s disease (AD), a cause of progressive cognitive decline, remains challenging. Recent information-theoretic advances allow brain dynamics to be quantified in terms of how regions share and combine information. Integrated Information Decomposition (ΦID) separates redundant (the same content present in multiple regions) from synergistic information (new content that emerges only when regions are considered together). Such information-dynamic measures may provide biomarkers relevant to AD risk and progression.

**Methods:** Here we applied integrated information decomposition (ΦID) to resting-state fMRI from the Alzheimer’s Disease Neuroimaging Initiative (ADNI), to test whether ΦID measures are diagnostically sensitive and track cognition along the AD spectrum. For each region, we computed total synergy and redundancy and compared values across cognitively normal (CN), mild cognitive impairment (MCI), and AD groups.

**Results:** Compared to CN, AD patients showed a striking synergy reduction across the entire brain, in concert with widespread redundancy increases, particularly in the executive and default mode networks. Transitions from CN to AD included an intermediate MCI decrease in redundancy, possibly reflecting early disease compensation strategies. This AD informational shift from complex higher level information processing to more robust inefficient forms likely reflects a cognitive shift to simpler, less integrative cognitive processes. Indeed, when re-analysing the data according to a standard cognitive clinical test (the Montreal Cognitive Assessment), we found a synergy-redundancy shift in high versus low performers broadly very similar to the CN to AD shift.

**Conclusion:** AD shows a clear information-processing signature: reduced global synergy and increased redundancy, especially in the executive control network. These striking results provide powerful insights into the widespread information processing reconfiguration that occurs in AD, with clear changes already emerging at the earlier MCI stage. Further, these results provide a novel route to support early diagnosis and stratification.

## 1 Introduction

Alzheimer’s disease (AD) is the leading cause of dementia and is associated with progressive cognitive decline and impairment in the activities of everyday life [48, 69]. AD imposes substantial personal and societal costs [9, 41], however it can be challenging to accurately diagnose at early or preclinical stages [3]. Biomarkers are increasingly used to support early diagnosis [30, 31], and new biomarkers should combine early detection with a mechanistic explanation for cognitive decline.

Information theory has the potential to serve these dual goals [60]. Beyond connectivity strength or linear coupling, information theory can characterise how brain regions exchange and combine information [20, 21, 62]. One tool of contemporary information theory, Integrated Information Decomposition (ΦID) [44], formally separates interactions into redundant (the same information present in multiple regions) and synergistic (information that emerges only when regions are considered together, i.e., not recoverable from parts alone) components. The information-theoretic approach has been applied to understanding many cognitive effects, both in healthy and in clinical populations, including some work exploring AD [8, 22, 39, 43, 45, 46, 53].

In the current work, we directly test the potential for ΦID [44], a method for exploring information dynamics, to provide mechanistic insight and diagnostic utility, using resting state fMRI data from the Alzheimer’s Disease Neuroimaging Initative (ADNI). ΦID has been applied to resting state data from the Human Connectome Project (HCP), where it was shown to be strongly related to both structure and function in healthy individuals [39, 64], and we apply here it to individuals with AD, those with mild cognitive impairment (MCI), and a cognitively normal (CN) group.

## 2 Methods

### Alzheimer’s Disease Neuroimaging Initiative

Data used in the preparation of this article were obtained from the Alzheimer’s Disease Neuroimaging Initiative (ADNI) database (adni.loni.usc.edu). The ADNI was launched in 2003 as a public-private partnership, led by Principal Investigator Michael W. Weiner, MD. The primary goal of ADNI has been to test whether serial magnetic resonance imaging (MRI), positron emission tomography (PET), other biological markers, and clinical and neuropsychological assessment can be combined to measure the progression of MCI and early AD.

Participants with at least one available 10-minute resting state fMRI scan prior to November 22nd 2023 were selected from the ADNI database, giving a total number of 2907 scans. The ADNI data collection has consisted of multiple phases, ADNI 1, ADNI GO, ADNI 2, ADNI 3 and ADNI 4. Of these acquisition rounds, resting state fMRI acquisition became standard from ADNI 3 onwards (2017) and remains part of the ADNI 4 protocol (2022).

One of the limitations of the ADNI data is that many participants entered the study in different phases, so fMRI scans are unavailable at baseline for a large portion of the dataset. For the purpose of our analysis, we opted to instead select the most recent scan acquisition available for each participant. Although many participants had several scans, potentially allowing for a longitudinal analysis, we opted in this case only to select one scan per participant, due to the aforementioned variation in scan procurement. A breakdown between the three diagnostic groups CN, MCI, and dementia, presumed AD, is given in the table below. We note, in particular, that a sizeable number of scans selected did not have diagnostic data available at the point of the scan and were excluded.

|  | No diagnostic data | CN | MCI | AD | Total |
| --- | --- | --- | --- | --- | --- |
| All scans | 179 | 1322 | 1053 | 353 | 2907 |
| Participants (scans selected) | 0 | 568 | 369 | 181 | 1118 |

### ADNI functional data acquisition

From ADNI2 onwards, a 10-minute resting state functional MRI acquisition was performed at every in-clinic visit. For ADNI2 this was solely on Philips scanners. For ADNI3 and ADNI4, all scanning was done on 3T MRI machines with two different protocols; basic (TR = 3 seconds for machines without multiband imaging) and advanced (TR = 0.6 seconds for machines with multi-echo capabilities). Participants were scanned with eyes open. For more information and individual scanner configurations see https://adni.loni.usc.edu/methods/documents/. For greater statistical power we used both the basic and advanced multiband images in our analysis, down-sampling the advanced protocol so that all sampling was done at an equivalent TR of 3 seconds.

### Conversion to BIDS format

We used Clinica’s ADNI-to-BIDS tool [56, 57] to convert the ADNI dataset to the Brain Imaging Data Structure [24], a standardised data structure for storing neuroimaging data and its derivatives. The BIDS structure was designed with the goal of increasing reproducibility within neuroimaging studies.

The raw and converted data were stored and processed on Queen Mary University of London’s High-Performance Computing cluster, Apocrita, supported by QMUL Research-IT http://doi.org/10.5281/zenodo.438045.

### Preprocessing with fMRIPrep

We used fMRIPrep 23.0.2 [16, 17], a tool based on Nipype [23, 25] to preprocess the anatomical and functional data for all groups. fMRIPrep provides a standardised pipeline for functional MRI processing using the BIDS data structure [24], and performs both anatomical and functional preprocessing steps. As a BIDS app, fMRIPrep is designed to make comparisons between studies more straightforward by standardising processing pipelines.

The anatomical preprocessing pipeline corrects field non-uniformity, strips the skull, segments and normalises onto MNI152Lin standard space with a resolution of 2mm. The functional pipeline estimates head motion (including three rotation and three translation confound parameters), before applying spatiotemporal filtering and slice-time correction, where possible. White matter, grey matter and CSF signals were extracted in addition to component-based noise features (tCompCor and aCompCor [4]) up to 50% of the variance. Finally, the functional data was resampled onto MNI152Lin standard space.

For full details on both the anatomical and functional preprocessing we refer to the boilerplate output from fMRIPrep given in the supplementary materials in appendix.

### Further denoising and cleaning steps

Following the work in healthy participants by Luppi et al. [39], we regressed out the first five anatomical CompCor [4] components for both cerebrospinal fluid and white matter, along with translations and rotations for all three dimensions and their derivatives as was made available through confound outputs from fMRIPrep. We also regressed out a non-steady state component provided by fMRIPrep. Also following the work by Luppi et al., we performed a band-pass filter between 0.05Hz and 0.0025Hz. No blurring step was performed so as to prevent “information leakage” between regions, which would falsely indicate shared information between nearby regions. As ADNI uses two main fMRI acquisition protocols (**basic** with a TR = 3 and **advanced** with TR = 0.6), we downsampled slices so that, to the nearest slice, all scans had a comparable TR of about 3s.

### Brain parcellations

We used the Schaefer-200 and Schaefer-100 parcellations [59] to parcellate the fMRI images into ROIs. These were supplemented with additional sub-cortical regions (32 and 16) respectively [61], to recreate the extended Schaefer atlases used on previous work with healthy participants [39], giving parcellations of size 232 and 116 respectively, which we refer to as ‘extended Schaefer’ atlases. These parcels were categorised according to the 7 networks of Yeo et al. [71], along with the subcortical regions as an eighth collection to give 8 networks/macroscopic regions, which we use repeatedly in our analyses.

### BOLD time series extraction

For each region of interest in the extended Schaefer-232 (116) atlas we used Matlab to select the corresponding voxels in each fMRI session and take an average at each time step across all voxels in the region, giving a new region-wise time series of BOLD signal.

### Partial Information Decomposition (PID)

Our analysis uses Integrated Information Decomposition (ΦID), a method built to extend an earlier method called Partial Information Decomposition (PID). The Partial Information Decomposition framework was proposed by Williams and Beer in 2010 [67] as a tool for understanding how information is stored in a system of variables. In particular, given a series of source variables *X*_1_, …, *X*_*n*_, which we assume we have knowledge of, how much information do we receive about a target variable *Y*, and how do the source variables qualitatively represent it?

It might be that several of the source variables provide the *same* information about *Y* (called **redundant** information), or that they give different, **unique** information about the target. Interestingly, it is also possible that knowledge of multiple sources can provide more information than any subset of the sources alone. This information is called **synergistic** information, and it reflects a kind of *additional deductive capability*, over and above any source used singularly. In this work, we focus primarily on ΦID’s generalisation of the **redundant** and **synergistic** information.

Knowledge of all sources together provides the maximum amount of information possible about *Y*, and this is the **mutual information**

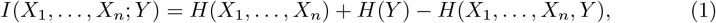

which quantifies the total amount of information in bits that *X*_1_, …, *X*_*n*_ provide about *Y* and vice versa [60]. As such, the contribution to knowledge of *Y* by all pieces of partial information (called **atoms**) should total the mutual information.

To compute these different atoms, a partial order is constructed on the information provided by sources. Information which is provided redundantly about *Y* by two collections of sources *A*_1_, …, *A*_*j*_ and *B*_1_, …, *B*_*k*_ is written

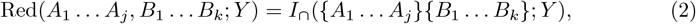

where the curly brackets signify information that is provided by all bracketed groups. The function *I*_∩_ is called a redundancy function or measure of intersection information: it quantifies how much information is shared by all bracketed groups about the target.

If a subset *S* ⊂ { *A*_1_, …, *A*_*j*_ } also shares information with *B*_1_, …, *B*_*k*_, then a partial order can be given by

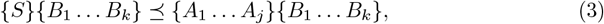

where the order ⪯ represents ‘informativeness’. That is, this shared information is inherited (in some sense) by a larger collection of variables. At each level, we expect that *more* information is made available about the target *Y* by adding more variables.

For this reason, the **change** as we go up the order ⪯ should reflect information which is exclusively provided at that level. For example, in a system of two source variables *X*_1_ and *X*_2_ and a target *Y*, the information uniquely provided by *X*_1_ is given by

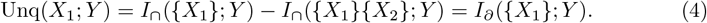

That is, information uniquely provided by *X*_1_ about *Y* is all the information provided by *X*_1_ about *Y* minus the information which is also given by *X*_2_ about *Y*. The notation *∂* is used to signify the direct contribution of information at this level in the order which is not available below it.

This process, isolating the individual atoms by subtracting those contributions lower in the order, is called **Möbius inversion** [13], and allows us to decompose information into non-intersecting pieces. It is the primary method used in information decomposition and set-theoretic perspectives on information theory [15, 44, 67].

At the top of the order is the **synergistic** atom, which reflects to what extent a collection of variables provides more information than the sum of its parts. For two variables this is given by

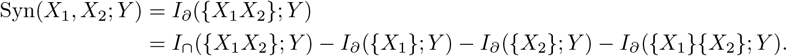

In our scenario, we decompose the joint evolution of a pair of regions 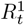 and 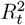 and their state after one time step (TR), 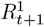 and 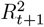 (recall that since the TR is 3 seconds, the variable observations are 3 seconds apart). That is, we decompose the **time-delayed mutual information**

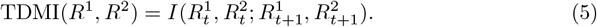

Although the structure of the order is provided by the partial information decomposition framework, the exact choice of how to calculate redundancy { *X*_1_ } { *X*_2_ } is hotly contested. Despite this, many partial information decomposition methods show similar behaviour for Gaussian systems— a PID referred to as minimum mutual information (MMI) [2, 39]. This PID sets redundant information as the lowest of the marginal contributions *I*(*X*_1_; *Y*) and *I*(*X*_2_; *Y*). Following Luppi et al., we also use this PID measure as brain activity is suitably Gaussian [5], following previous work in neural information processing. In the supplementary, we also revalidate our results using a ΦID approach due to Ince based on common change in surprisal [29].

For two source variables and one target variable, partial information gives rise to 4 new information quantities over and above classical information theory: the redundant information (shared between the sources about the target), two unique components (unique to each variable), and a synergistic component (available only by observing both source variables at the same time). These four pieces are further refined in integrated information decomposition (ΦID) before being used in our analyses.

We note briefly that other decompositions of information also exist with potential to characterise synergy and redundancy [14, 15, 55], along with some non-decomposition methods such as the *O*-information [54]. For this work, we used the ΦID method, an extension of PID, to describe synergy and redundancy, although other approaches are viable.

### Integrated Information Decomposition (ΦID)

The partial information decomposition approach in its current form only allows for a single target, and does not consider how information might be stored across multiple target variables. For this reason, Mediano et al. [44] introduced **Integrated Information Decomposition** (ΦID), to explore how information provided by the sources is stored by multiple target variables, hence broadening the *taxonomy of information dynamics*.

Integrated information decomposition can be thought of as describing a *forwards* and *backwards* partial information decomposition at the same time. To see how this works, consider two regions of interest *R*^1^ and *R*^2^. We have the forward PID redundancy (information in the present which is stored redundantly about the future),

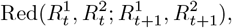

and we have the reverse PID redundancy (information in the present which is stored redundantly about the past),

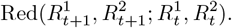

ΦID posits that we can reasonably ask how much information stored redundantly at time step *t* was also then stored redundantly at time step *t* + 1, giving a double-redundancy atom 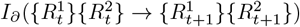. For each combination of PID atom in the present and PID atom in the future, there is a corresponding ΦID atom, meaning that 16 such combinations are possible for a system of two targets and two sources. All together, providing a PID redundancy function and the ΦID double redundancy function is enough to specify all 16 atoms in the decomposition.

In this work, unless otherwise qualified, we refer to 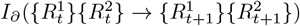 as **redundancy** and 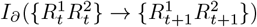 as **synergy** without further qualification. For later analyses based on the taxonomic decomposition introduced in the original ΦID paper, we use *r* to represent redundancy, *x* and *y* to represent information unique to the first and second region respectively, and *s* to represent synergistic information. We will occasionally make use of the notation *r* → *r* or *rtr* to designate information that was redundant in the past and in the future. For example, we write *rts* to represent information stored redundantly in the past which is stored synergistically in the future. We mostly focus on the redundancy and synergy atoms in this work as per Luppi et al.’s approach, and which can be used to characterise an information-functional hierarchy [39].

While ΦID-MMI guarantees the positivity of the redundancy and synergy atoms, many of the other atoms can occasionally take negative values. Some authors interpret this as an embedded misinformation behaviour, or representing a sort of natural mathematical dual to redundancy (shared ignorance) [18]. In analyses where these atoms appear we examine relative changes in these atoms between diagnostic or cognitive groups, rather than their absolute values. In other results we focus on the non-negative synergy and redundancy terms.

### Matrix of redundant and synergistic interactions

Following previous work by Luppi et al. [39] on data from the Human Connectome Project (HCP), all of the sixteen ΦID atoms were computed for every possible pair of ROIs for every participant for the extended 232 Schaefer atlas (also replicated for the extended 116 parcellation). Following Luppi et al., we decomposed the TDMI with ΦID between every pair of regions 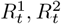 and their state in the next echo 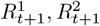,, giving a matrix of information dynamics for each scan. These matrices are provided for the sts and rtr atoms in the appendix in section 6.7.

For results on diagnostic state we averaged these matrices across all participants in each diagnostic group — cognitively healthy participants, participants with MCI, and those with dementia at the time of the scan, giving three 232 × 232 (116 × 116) matrices depicting *average* values between each pair of ROIs for each atom (rtr, rtx, rty, rts, xtr, xtx, xty, xts, ytr, ytx, yty, yts, str, stx, sty, sts).

This gives a 232 × 232 matrix of information dynamics between pairs of regions, representative of the average information dynamics across each diagnostic group for a given atom. We then cleaned up these matrices by regressing out age, years in education, gender, and intracranial volume (available either from the ADNI dataset, or where ICV was not available, we imputed it using data generated by fMRIPrep).

The precise behaviour of the null distribution for ΦID atoms is difficult to determine in this case; although permutation testing would be optimal (e.g. in a manner similar to the approach taken in [37]), due to the large quantity of data, this was mostly intractable for our calculations. For that reason, we tested all pairwise changes between the CN and AD groups in the redundancy and synergy atoms using independent two-tailed *t*-tests at the *α* = 0.05 level, accounting for multiple comparisons (232 × 232 × 16 = 861, 184) using the Benjamini-Hochberg procedure for constraining the false discovery rate. This result is reported in the supplementary materials in section 6.7. Significant increases and decreases were separated from each other prior to plotting. Regions were also re-ordered so that regions in each of the 7 Yeo networks (and subcortical regions) are grouped together [71].

### Reduction to a regional measure

Following Luppi et al. [39], for each pair of regions of interest (ROIs) in the Schaefer parcellations, we computed all of the ΦID atoms and obtained an average information dynamics matrix for each participant group as above. In results 3.1, 6.3, and 6.10, we took the sum across each column of this matrix to reduce these pairwise interactions to interpretable behaviour at a regional level. For example, to compute the rtr for a given region *R*^*i*^, we summed down the matrix column:

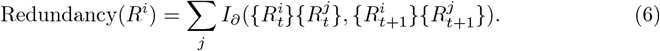

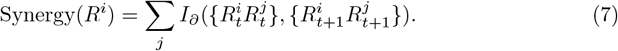

All self-measures (*R*^*i*^, *R*^*i*^) were taken to be zero. Comparisons were done regionwise using an FDR-corrected Welch’s *t*-test.

### Intranetwork group comparisons

In addition to comparing the total summed redundancy and synergy for each region with all other regions, we also performed a network-wise analysis by averaging all intranetwork redundancy and synergy within each network. For regions *i* in some set of regions corresponding to a network *S*, we computed

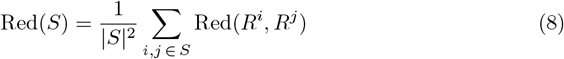

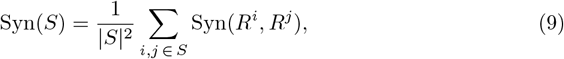

before using FDR-corrected Welch’s *t*-tests to detect statistically significant ΔRed and ΔSyn at the *α* = 0.05 significance level.

### Regional *z*-scoring against the CN group

For the qualitative depiction in result 3.1, we selected the average summed synergy and redundancy atoms from each region across the MCI and AD groups (repeating in the supplementary with the Medium MoCA and Low MoCA), before *z*-scoring them against that region’s summed measures across all scans in the CN group (or the High MoCA group, respectively). That is to say, each region’s redundant and synergistic activity in MCI/AD (Medium/Low MoCA) is scored based on how unusual its value would be in the CN distribution. These *z*-scores were then plotted against each other in result 3.1 to show a qualitative picture of changes in information dynamics.

Note this is done region-wise. For each MCI/AD region, the CN mean for the summed atom of that region is subtracted to give a mean of zero, before being divided by the deviation for that region to normalise onto a *z*-score.

### Projection onto the first principal component

In result 3.1 we also project synergy and redundancy onto the first axis of variation. This was done in python by projecting the variation in synergy and redundancy onto its first component using NumPy. Regions were coloured on this plot based on their location in the 7 Yeo networks (or if that region was a supplementary subcortical region). The variance explained in each diagnostic (cognitive) group by PC1 was also extracted.

### Cortical plotting

Cortical plots were created using the ENIGMA toolbox’s `plot_cortical` function in Matlab with custom colour palettes [36]. In figures 1 a) and 4 a), the cortical plot is the first principal component of *z*-scored changes to redundancy and synergy. In figure 1 b) and 4 b), the *z*-scores are used in redundancy and synergy, respectively separated.

**Figure 1.**
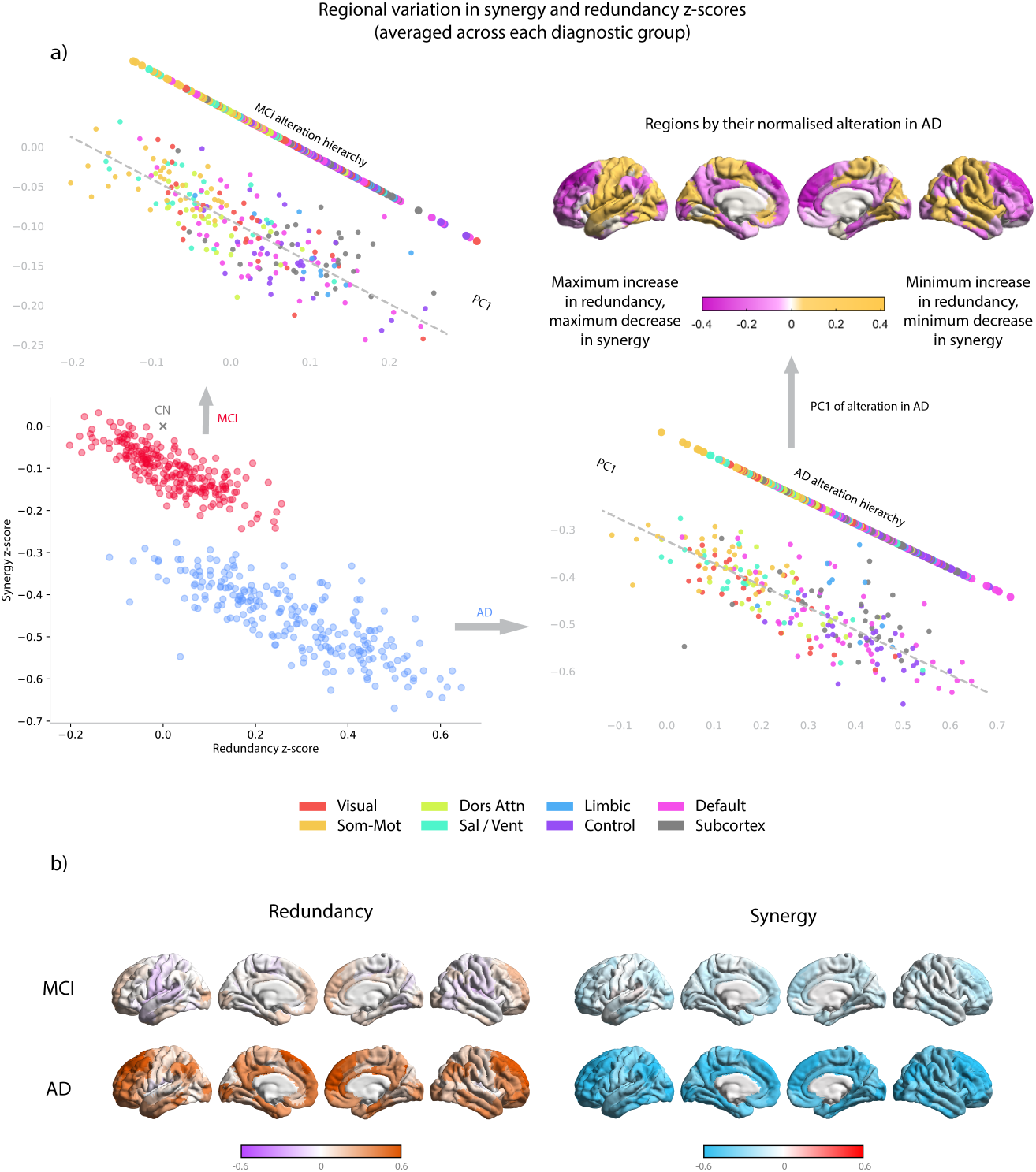
Information dynamics are altered according to functional roles. Figure a) Bottom left: 232 regions from the extended Schaefer parcellation averaged across all participants labelled as having MCI (red) and dementia (blue). Regions are z-scored against the CN mean (grey cross) for each region in their total amount of redundancy and synergy. Top left: z-scores for regions from the MCI group, with Yeo networks labelled (subcortical regions depicted in grey). This is then projected onto the first principal component of variation across all 232 regions to show that changes in information dynamics are modulated by the unimodal-transmodal axis. Bottom right: as with the top left, but with the AD group. Top right: each region is plotted onto the cortex with its first principal component (PC1) scores from the AD *z*-scores. Regions with maximum increases in redundancy and maximum decreases in synergy are in pink, minimum increases in redundancy and minimum decreases in synergy are in yellow. Figure b) Redundancy and synergy *z*-scores given separately. Left: regions with increasing redundancy are given in orange; regions with decreasing redundancy are given in pink, and increases in orange. Right: regions decreasing in synergy are highlighted in blue; increases are depicted in red.

For result 3.1 and figures 2 and 9 we used plots from [43], originally constructed with Brainnetviewer [70], with some recolouring.

**Figure 2.**
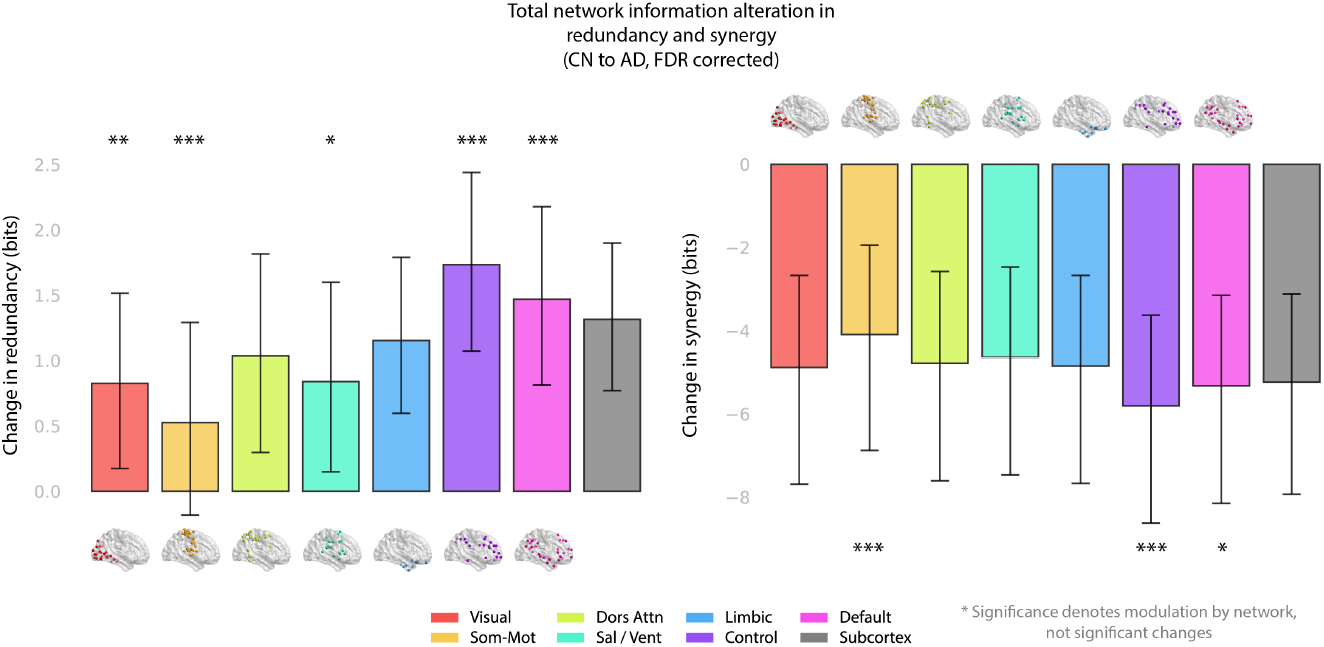
Information changes from CN to AD are heterogeneous across brain networks. For each region the total redundancy and synergy with all other brain regions is calculated, before being averaged across each network to find the average total increase or decrease in that network’s average regional information processing. Left: increases in total redundancy averaged across each of the Yeo networks (plus subcortex); right: decreases in total synergy averaged across each of the Yeo networks. Significant deviations from a global average were computed using a permutation test with *n* = 10, 000 resamples and FDR correction to account for multiple comparisons, shuffling network labels rather than diagnostic labels to test the hypothesis that changes in information dynamics (redundancy or synergy) are modulated by region, not solely by a global effect. Significances are illustrated at the *α* = 0.05^∗^, 0.01^∗∗^, and 0.001^∗∗∗^ levels respectively. Confidence intervals were computed by bootstrap resampling to obtain the 95% confidence interval.

### Verifying functional modulation of information processing changes

In result 3.1 and figures 2 and 9 we averaged the summed measure across each of the 8 macroscopic network regions (7 Yeo networks, plus 32 subcortical ROIs). To test whether function (and not a whole-brain effect) modulated the synergistic decrease and redundant increase when comparing CN to AD, we used a permutation test with *n* = 10, 000 resamples, shuffling network labels. An FDR correction was also applied for the 8 tests performed for synergy and redundancy respectively.

### Taxonomic information signature calculation

The taxonomic calculation in result 3.3 used the intranetwork values for individual ΦID atoms, grouped together in a fashion similar to the modes (taxa) described in the paper introducing ΦID [44]. Depending on the qualitative behaviour of the individual atoms, they can be grouped into 6 broad modes:

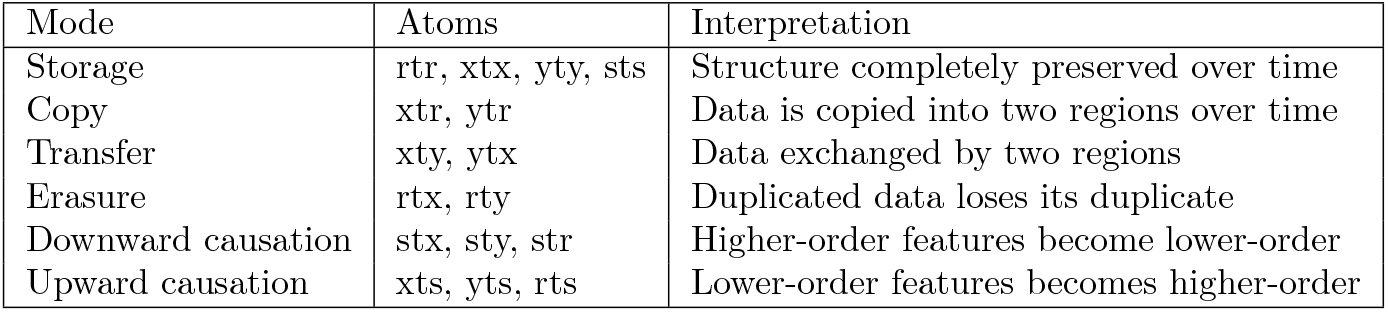

| Mode | Atoms | Interpretation |
| --- | --- | --- |
| Storage | rtr, xtx, yty, sts | Structure completely preserved over time |
| Copy | xtr, ytr | Data is copied into two regions over time |
| Transfer | xty, ytx | Data exchanged by two regions |
| Erasure | rtx, rty | Duplicated data loses its duplicate |
| Downward causation | stx, sty, str | Higher-order features become lower-order |
| Upward causation | xts, yts, rts | Lower-order features becomes higher-order |

In principle these quantities can all vary independently of each other, but due to additional constraints imposed by the MMI-ΦID, the minimum unique information in either direction must be precisely zero by construction. This gives

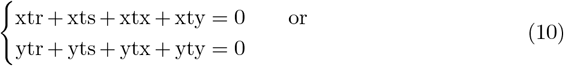

and

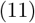

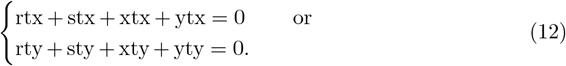

Overall, for any pair of signals *R*^1^ and *R*^2^, 8 of the 16 ΦID atoms calculated between those two signals with MMI-PID will, assuming non-negativity, be zero. In practice this is frustrated by negativity considerations, but the total number of degrees of freedom is certainly reduced.

In empirical practice we found that the copy and transfer atoms (dual to each other in direction) and the upwards and downwards causation modes (also dual) behaved structurally very similarly. For this reason, we grouped into four dominant modes, **storage, transfer, copy/erasure**, and **multi-scale causation**.

By summing up the corresponding atoms, we computed intranetwork values for each mode in each diagnostic (MoCA) group. For each mode-network pair, we tested for modulation across groups using a one-way ANOVA with an FDR correction across all 32 tests, reporting at the *α* = 0.05, 0.01, 0.001 levels. The same test was applied to MoCA categories, with results given in the supplementary.

Post-hoc tests were performed on pairs of groups (pairs of diagnostic groups or MoCA categories) for each one-way ANOVA which was significant, using Welch’s *t*-test to compare means without assumptions of variances, adjusted for the false discovery rate at the *α* = 0.05 level. Significant differences are reported in tables 5 and 6.

## 3 Results

### 3.1 A strong principal axis captures heterogenous synergy and redundancy changes to ROIs in AD

#### 3.1.1 Organisation of changes in summed information dynamics across diagnoses

In order to qualitatively depict how redundancy and synergy are changed in different ROIs and networks when comparing CN, MCI, and AD groups, we computed the mean synergy and redundancy, along with their standard deviations, for each ROI across the CN group. Using these values, we *z*-scored the average synergy and redundancy for each ROI from the MCI and AD groups, giving an indication of how each region changed relative to a cognitively healthy baseline. A corresponding plot of these scores can be seen in figure 1, where each dot represents an ROI from either the average MCI information dynamics or average AD information dynamics. Network-wise *z*-scores are available in the supplementary materials.

In figure 1 a) we show the regional change in information processing dynamics between the CN group and the MCI and AD groups. For each region *i*, the synergistic information and redundant information are summed across all interactions with all other regions *j*, giving a single measure of a region’s synergistic and redundant behaviour. For scans where the individual is labelled as CN, the mean and standard deviation was computed for each region’s redundancy and synergy totals. The average regional synergy and redundancy in all MCI (red) and AD (blue) acquisitions was found and *z*-scored against the CN mean and standard deviation for that region. The corresponding scores are plotted against each other in the bottom left. In order to better see the function of each region, the top left and bottom right panels show the same regions, now coloured with each of the Yeo networks [71] (plus an additional subcortical grouping [61]), along with a projection onto the first principle component (PC) of variation. In the top right of 1 a) there is a cortical plot of the AD group projected onto the first principal component (PC1), showing that greatest regional change is observed in the limbic, ECN, and DMN systems. Figure 1 b) shows the synergy and redundancy *z*-scores independently of each other, plotted onto the cortex.

When arranging brain regions in this way (changes in redundancy and synergy), variation occurs mostly across a single axis, with the first principal component capturing 92.3% of the regional variation in MCI and 93.2% of variation in AD.

We repeated this procedure with three categories based on the Montreal Cognitive Assessment (MoCA) [47], with results pointing to a similar structure. The corresponding results are available in the supplementary materials in section 6.1.2.

#### 3.1.2 Changes between CN and AD

When looking at individual ROIs, we applied Welch’s *t*-test to each of the 232 parcels and applied an FDR correction to test for significant differences between the CN and AD groups. At the *α* = 0.05 level, all 232 regions from all networks exhibited a significant decrease in synergy. In redundancy, a majority of executive control network (ECN) regions (29/30), default mode network (DMN) regions (39/46), subcortical regions (27/32), and limbic regions (10/12) exhibited a significant increase in the AD group compared to the CN group. Significant increases in the same comparison were also observed in the dorsal attention network (11/26 ROIs), visual system (10/29 ROIs), salience network (6/22 ROIs), and somatomotor network (3/35 ROIs), but with lower coverage of the network seeing a significant increase.

In addition to the individual ROIs, we also performed analyses on the intranetwork average (looking at pairwise interactions inside of each network). We observed a significant decrease in the AD group compared to the CN group in redundant interactions in the somatomotor network (ΔRed = − 0.009 bits, FDR-adjusted *q* = 0.004, Hedges’ *g* = − 0.242), and in synergistic interactions in the visual (ΔSyn = − 0.018 bits, *q* = 0.008, *g* = − 0.290), dorsal attention (ΔSyn = − 0.016 bits, *q* = 0.013, *g* = − 0.267), salience (ΔSyn = − 0.012 bits, *q* = 0.046, *g* = − 0.215), limbic (ΔSyn = − 0.019 bits, *q* = − 0.003, *g* = − 0.349), control (ΔSyn = − 0.026 bits, *q* < 0.001, *g* = − 0.498), and default mode networks (ΔSyn = − 0.020 bits, *q* = 0.002, *g* = − 0.387), along with the subcortex (ΔSyn = − 0.023 bits, *q* < 0.001, *g* = − 0.446). Statistically significant increases in the AD group compared to the CN group were found in redundancy in the limbic (ΔRed = 0.005 bits, *q* = 0.036, *g* = 0.214), control (ΔRed = 0.008 bits, *q* < 0.001, *g* = 0.416), and default mode networks (ΔRed = 0.004 bits, *q* = 0.030, *g* = 0.223), as well as in the subcortex (ΔRed = 0.005 bits, *q* = 0.008, *g* = 0.280).

#### 3.1.3 Changes between CN and MCI

Before applying an FDR correction, (5/32) regions in the subcortex, (6/32) in the default mode network, (3/39) in the executive control network, (1/12) in the limbic network, and (1/29) in the visual network exhibited significant increases in redundancy in MCI when compared to the CN. Similarly, (8/12) regions in the limbic system, (15/32) in the subcortex, (14/30) in the ECN, (20/46) in the DMN, (5/29) in the visual network, (4/26) in the dorsal attention network, and (1/22) regions in the salience network showed decreased synergy in MCI when compared to the CN group, but none of these results (redundancy or synergy) survived the FDR correction.

When performing an analysis on intranetwork averages inside of each of the 8 networks, we observed only one significant difference surviving FDR correction, which was a decrease in redundancy in the somatomotor network in the MCI group when compared to CN individuals (ΔRed = − 0.009 bits, *q* < 0.001, *g* = − 0.269). Prior to FDR correction, a decrease was also detected in redundancy in the salience network along with an increase in synergy in the somatomotor network, but these did not survive the multiple-comparisons correction.

#### 3.1.4 Changes between MCI and AD

When comparing the MCI to the AD group with an FDR correction, we saw an increase in redundancy in AD in (23/30) regions in the ECN, (35/46) in the DMN, (13/26) in the dorsal attention network, (15/32) in the subcortex, (9/22) in the salience network, (8/35) in the somatomotor network, (6/29) in the visual network, and (2/12) in the limbic system. Similarly, we saw significant decreases in synergy in AD in all regions (30/30) in the ECN, (34/35) in the somatomotor network, (25/26) in the dorsal attention network, (21/22) in the salience network, (29/32) in the subcortex, (40/46) in the default mode network, (25/29) in the visual system, and (9/12) in the limbic system. No increases in synergy or decreases in redundancy survived FDR correction.

For the intranetwork average comparison, we found significant increases in redundancy in AD compared to MCI in the executive control network (ΔRed = 0.008 bits, *q* = 0.005, *g* = 0.344), default mode network (ΔRed = 0.005 bits, *p* = 0.021, *g* = 0.269), and subcortex (ΔRed = 0.004 bits, *q* = 0.043, *g* = 0.213), while we observed significant decreases in synergy in AD compared to MCI in the visual (ΔSyn = −0.016 bits, *q* = 0.035, *g* = −0.251), dorsal attention (ΔSyn = −0.015 bits, *q* = 0.035, *g* = −0.242), limbic (ΔSyn = −0.014 bits, *q* = 0.035, *g* = −0.245), control (ΔSyn = −0.020 bits, *q* = 0.005, *g* = −0.361), default mode networks (ΔSyn = −0.017 bits, *q* = 0.016, *g* = −0.311), along with the subcortex (ΔSyn = −0.017 bits, *q* = 0.014, *g* = −0.316).

### 3.2 Synergy loss and redundancy gain in AD are not uniform across networks

All *p*-values in this section are given with FDR correction. Taking averages across all of the 7 Yeo networks plus the subcortical collection of atoms, significant decreases in total synergy between the CN group and the AD group were observed in all 8 brain networks with effect sizes largest in the executive control network (ECN) (Welch’s *t* = −4.509, FDR-adjusted *q* < 0.001, Hedges’ *g* = −0.510), subcortex (*t* = −4.215, *p* < 0.001, *g* = −0.475) and default mode networks (DMN) (*t* = −4.135, *q* < 0.001, *g* = −0.467). There were still moderate effect sizes in the limbic (*t* = −3.785, *q* < 0.001, *g* = −0.433), visual (*t* = −3.731, *q* < 0.001, *g* = −0.415), dorsal attention (*t* = −3.666, *q* < 0.001, *g* = −0.402), and salience networks (*t* = −3.610, *q* < 0.001, *g* = −0.397). The smallest change was seen in the somatomotor network, although this difference was still strongly significant (*t* = − 3.220, *q* = 0.001, *g* = − 0.349).

When looking at total redundancy across each Yeo network, the ECN once again showed the strongest effect when comparing CN to AD (Welch’s *t* = 4.948, FDR-adjusted *q* < 0.001, Hedges’ *g* = 0.491), followed by the subcortical (*t* = 4.484, *q* < 0.001, *g* = 0.448), default mode (*t* = 4.169, *q* < 0.001, *g* = 0.419), and limbic (*t* = 3.707, *q* < 0.001, *g* = 0.408) networks. Unlike for synergy, there is then a strong drop off in effect size. Despite showing a smaller overall effect, changes were still significant in the dorsal attention (*t* = 2.668, *q* = 0.013, *g* = 0.244), visual (*t* = 2.408, *p* = 0.022, *g* = 0.225), and salience (*t* = 2.261, *q* = 0.028, *g* = 0.209) networks. The changes to redundancy in AD compared to CN did not reach the threshold for significance in the somatomotor network (*t* = 1.379, *q* = 0.169, *g* = 0.124).

Changes in synergy and redundancy for each network, along with a different statistical test for network-specificity, are shown in figure 2, with error bars found using bootstrap resampling for a 95% confidence interval.

#### 3.2.1 Network modulation is not simply due to a global effect

To explore the idea that increases in redundancy and decreases in synergy are inhomogeneous across brain **networks**, we tested the hypothesis that individual Yeo networks exhibited their own information dynamics in each group (rather than a global effect across all brain regions) using a permutation test, shuffling synergy and redundancy changes across ROIs. Results are shown for the 7 Yeo networks plus an additional collection of subcortical regions in figure 2.

When comparing the CN group to the AD group, the change observed is more homogeneous across brain regions. Network changes in synergy which could not be explained by a global effect appeared in the ECN (ΔSyn = − 5.806, FDR-adjusted *q* < 0.001), the DMN (ΔSyn = − 5.324, FDR-adjusted *q* = 0.009), and the somatomotor network (ΔSyn = − 4.092, FDR-adjusted *q* < 0.001).

Looking at the difference in redundancy, we saw a more heterogenous change across brain regions when comparing the CN group to those with AD. Once again the executive control network (ECN) saw the largest increase (ΔRed = 1.733, FDR-adjusted *q* < 0.001), followed by the default mode network (DMN) (ΔRed = 1.469, *q* < 0.001). Also significantly deviating from a global shift were the salience (ΔRed = 0.839, *q* = 0.032), visual (ΔRed = 0.826, *q* = 0.008), and somatomotor networks (ΔRed = 0.525, *q* < 0.001), which saw a smaller increase than the global network mean.

### 3.3 Changes in taxonomic information signatures in progression to AD and cognitive decline

One of the main strengths of ΦID is that atoms can be grouped together into larger *modes* which describe different classes of dynamic information processing. In order to better understand the mechanisms underlying the observed increase in redundancy and decrease in synergy, we used groups of atoms from the paper introducing ΦID to explore the information dynamics observed [44]. Due to the assumed stationarity of the distribution, we have simplified the original modes to four: *multi-scale causation, copy and erasure, transfer*, and *storage*.

The *multi-scale causation* mode reflects the amount of higher-order information representations and lower-order representations which are exchanged. In terms of cognition, we believe this to be directly related to the stability of different information representations in the brain. Where transitions into and out of higher-order representations are frequent, long chains of deductive information representations are less common.

The *copy and erasure* mode reflects where information is both increasingly duplicated and more often deleted^1^. In terms of cognition and brain dynamics, we expect this might correspond to a more macroscopic propagation of information and a delocalisation of specific information transmission; complexity in representations likely decreases.

The *transfer* mode describes information that was relocated from one region to another, without duplication. In terms of brain dynamics, this corresponds to a ‘clean information flow’ from region to region. Reduction in transfer likely represents difficulties with neuronal signal transmission, with signal often being dropped.

Finally, the *storage* mode represents information that remained in the same atom between the two time steps. In terms of neural representations, this corresponds to information that is persistently and robustly encoded, and not dropped or otherwise interfered with.

For each of the 7 Yeo networks (along with the additional subcortical areas), atoms inside of each mode (multi-scale causation, copy and erasure, transfer, and storage) were added together and averaged across all of the ***intranetwork*** interactions for those regions. For example, the *transfer* mode in the *limbic system* was found by averaging all of the xty and ytx (transfer) atoms between all pairs of regions within the limbic system and adding them together. The results, given here with an FDR-corrected one-way ANOVA, are shown in figure 3.

**Figure 3.**
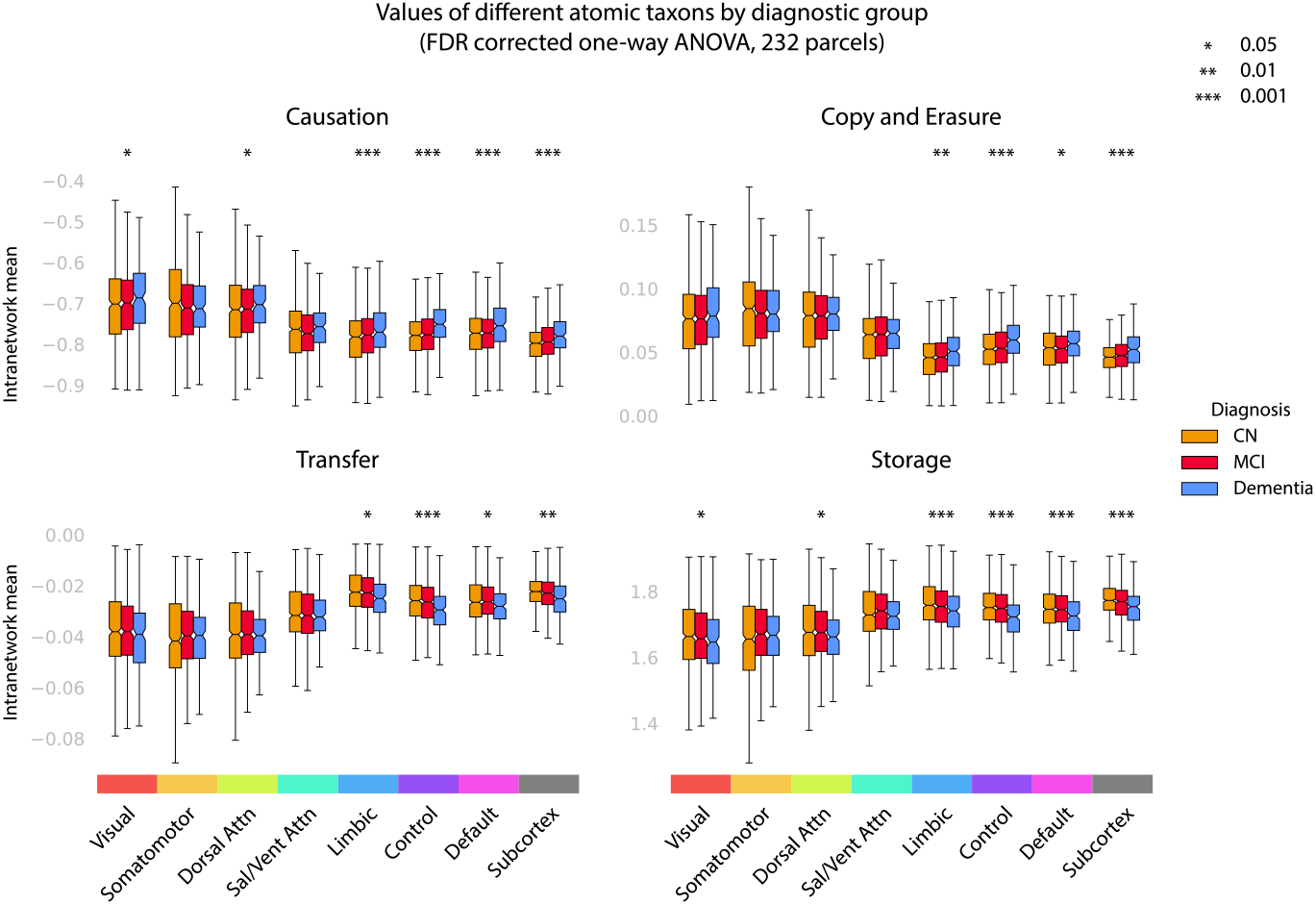
Taxonomic information processing is strongly modulated by diagnostic group in limbic, control, default mode and subcortical regions, with strong decreases in transfer and storage in AD. Four macroscopic modes (multiscale causation, copy and erasure, transfer, and storage) were averaged across all pairs of brain regions inside of 7 Yeo networks using the ΦID approach, along with a collection of subcortical regions. The upwards and downwards causation modes are grouped together into a **multi-scale causation** mode, and the copy and erasure modes are also grouped due to linear constraints. For each mode-network pair, we performed a one-way ANOVA, correcting for the FDR.

Across diagnostic groups, the multi-scale causation and storage modes saw the most significant results, with the omnibus ANOVAs locating a statistically significant group variation at the *p* < 0.001 level in the ECN, DMN, subcortex, and the limbic system, following a similar pattern as previously observed in sections 3.1 and 3.2. In addition to this, in the multi-scale causation and storage modes, the visual and dorsal attention networks varied across groups at the *p* < 0.05 significance level. In none of the four modes did the ANOVAs detect significant variation across groups in the somatomotor or salience networks.

Post-hoc Welch’s *t*-tests with FDR correction revealed that all of the group differences were driven by comparisons with the AD group, with no comparisons between the CN and MCI groups reaching the threshold for statistical significance. In the multi-scale causation and copy-erasure modes, significant increases were detected when comparing the AD group to the CN group, while in the transfer and storage modes, significant decreases were detected when comparing the AD group to the CN group.

The largest effects appeared when comparing CN to AD in the executive control network, which reached the *p* < 0.001 threshold for significance in all four modes (increasing with AD in multi-scale causation and copy-erasure, decreasing in transfer and storage). Full results for the post-hoc Welch’s *t*-tests are given in supplementary in table 5.

#### 3.3.1 Correspondence between information signatures in cognitive examination and diagnosis

For all of the results available here, similar patterns emerged when using three groups (high, medium, and low cognition) based on the Montreal Cognitive Assessment (MoCA) [47], and are presented in the supplementary materials 6. Most notably, broad global decreases in synergy were seen in the low MoCA group when compared to the high MoCA group, along with broad increases in redundancy, although the spread of synergistic change was greater than when comparing diagnostic categories.

As in the diagnostic analysis, significant decreases in total synergy were found to be highly present in the ECN, salience network, DMN, subcortex, dorsal attention network, and somatomotor network, with the fewest ROIs showing reduction of synergy in low MoCA in comparison to high MoCA in the limbic system. Similarly, increases in redundancy from high to low MoCA were seen heavily in the ECN, subcortex, and DMN, with some increases also seen in the dorsal attention network, salience network, limbic, and visual systems. Very few ROIs in the somatomotor network showed similar increases with cognitive decline.

Finally, as with the diagnostic categories, changes in redundancy and synergy between high and low MoCA cognitive scores were seen to highly depend on brain network and could not be explained solely by a global effect. All results on the MoCA categories are reported in full in the supplementary materials 6.

## 4 Discussion

In this work we applied recent advances in the theory of information decomposition to study how the brain represents information in AD and MCI using the large ADNI fMRI dataset, exploring two main information-theoretic effects: synergy, where different regions cooperatively integrate information, and redundancy, where regions represent duplicate information. The AD group exhibited major decreases in synergy (in all regions) and increases in redundancy (in a majority of regions) when compared to the CN group, with similar prodromal patterns already appearing in the MCI group (although this effect did not survive FDR correction). Moreover, we found that regional changes observed fell strongly onto a principal axis, ranging from highly-associative regions in the ECN and DMN (which saw the greatest decreases in synergy and greatest increases in redundancy) to less associative networks (such as the somatomotor network), which saw smaller effects. More than this, we showed that all our results are similarly explained by cognitive deficits using categories derived from MoCA scores [47], directly linking information dynamics to cognition. Overall, this work strongly demonstrates the efficacy of the information-dynamical approach to understanding cognitive deterioration in AD, and opens new avenues to the construction of information-theoretic biomarkers of cognitive decline.

Most of our results are best explained by connectonomic disruption, especially where it corresponds to AD-related atrophy (for example, in a Braak-like staging pattern [6, 7]). Synergistic effects are often fragile [51], while redundant information, which exhibits greater informational stability, might survive the neuronal disruption associated with amyloid and tau deposition, or possibly be more heavily utilised in order to overcome information loss associated with neuronal dysfunction. Our finding that multi-scale causation and copy-erasure modes are increased in AD further supports this hypothesis, along with the destruction of the transfer and storage modes in cognitive decline, which further highlight the possibility of signal disruption.

These results have implications for how we link high-level cognition to lower-level qualitative information processing mechanisms, opening up new avenues for exploring early diagnosis of major neurocognitive disorders with imaging techniques. Synergy, in particular, signifies a kind of information-theoretic *deduction*, so the especially strong decrease in synergy in the executive control network is insightfully consistent: as it has been shown that higher synergy is associated with highly-associative brain regions employed in complex cognition in healthy individuals [39], the decrease in synergy observed here might correspond to a decreased ability to synthesise and integrate information from multiple sources. Such a decline could render complex cognition more challenging, and this effect might be visible even at the MCI stage with careful biomarker development. Since synergy is often represented as a measure of emergent behaviour, and is also seen to be reduced in loss of consciousness [38, 39, 40], this work also gives further credence to the classification of AD as a disorder of consciousness (DoC) [28].

In addition to the broad increased redundancy and decreased synergy effect, we also found that changes in information dynamics in MCI and AD (and across our MoCA categories) were modulated by Yeo network [61, 71], with somatomotor regions demonstrating the greatest resistance to changes in information dynamics in AD compared to CN. In the AD group, the greatest changes were seen in the executive control and default mode networks, with subcortical regions also strongly affected. This pattern roughly approximates a Braak-like staging pattern [6, 7]. In this pattern, early tau deposition is observed in limbic and subcortical regions before an isocortical presentation at later stages (though somatomotor regions remain least affected). This structural decline can also be seen in our results: in the MCI group, we note that, despite not surviving FDR correction, the limbic system, subcortex, and ECN showed the strongest changes when compared to the CN group. This connection supports the hypothesis that the cellular transport disruption associated with tau deposition is implicated in changes in information representation— an intriguing mechanistic insight warranting further investigation.

To explore the qualitative changes in information processing with a more mechanistic lens we also explored intranetwork information dynamics by grouping atoms into ΦID modes: multi-scale causation, copy-erasure, transfer, and storage. We saw increases in the multi-scale causation and copy-erasure modes and decreases in the transfer and storage modes when comparing the AD group with the CN group, with the strongest changes seen in the limbic system, ECN, DMN, and subcortex, with somatomotor and salience networks least affected. Decreases in the transfer and storage modes are strongly supportive of the idea that information transmission and persistence is damaged in AD, indicating a direct disruption of information flow and decreased signal persistency. In terms of behaviour, we saw that this could potentially correspond to many cognitive difficulties: if information flow around the brain is interrupted, it could be linked to forgetfulness, or difficulties following chain-of-thought reasoning. Cognitive representations that would normally be stable and persistent (such as when focussing or holding attention), could inadvertently be dropped if local information storage declines. More work using qualitative information theory could illuminate the relationship to specific cognitive deficits in dementia.

In addition to the decreased transfer and storage, the observed increase in multi-scale causation might be due to a reduced stability of representations; if information cannot be stored persistently in the same fashion, alternative representation mechanisms might need to be more frequently substituted in, adding to this volatility. Neural pathways might become noisier, potentially swapping in and out of deductive cognition to understand and represent mental calculations, objects, and experiences. Increases in copy-erasure dynamics might represent an increase in *transient* data representations, or, perhaps as a counteraction mechanism, an attempt to *flood* a signal through a disrupted connectome, even if this signal lacks persistence. Representations might be split across multiple regions, providing more widely experienced (across areas), but more easily forgotten, cognition, a potentially jarring combination. Exploring these dynamics and their relationship to macroscopic behaviour and cognition will prove fruitful for understanding qualitative differences in brain activity in those with AD and cognitive impairment, along with the first-person experiences of such diseases.

One limitation of the ADNI fMRI dataset is that acquisitions are resting state, with participants not actively engaged in any cognitive task. This means that the signature we have obtained is subject to a large amount of noise: at any time, participants might be wind-wandering, recalling memories, or otherwise transitioning between different mental states. Because of this, the results we report here are only significant at the group level and do not, without further analysis, offer individual diagnostic strength. We expect that future work on task-based paradigms in cognitive impairment might offer a more participant-specific signature of information processing (see [43], for example). Given our results, we expect that tasks targeting the ECN, DMN, subcortex, or limbic system might offer the most potential for the development of an information decomposition-based diagnostic tool.

An additional limitation in this work is ongoing mathematical speculation about partial and integrated information decomposition methods. Many versions of both decompositions exist (see, for example, Kolchinsky’s commentary [34]), and precise choices for how to define redundancy have not yet been fully evaluated in the literature. The current view is that redundancy, as a concept, can be defined in multiple ways, and we make no particular philosophical commitments in this work. Despite this, we did validate our analyses with two different ΦID approaches, with results available (showing consistent patterns to those found in the main body) in the supplementary materials in section 6.13.

Similar work has appeared in the literature due to Zhang et al. [72], however their analysis focusses primarily on the graph-theoretic properties of synergy and redundancy. Our analyses focus on atomic differences themselves. In particular, we explore how these measures change between CN individuals, those with MCI, and those with AD. In the supplementary materials we also explore three different categories of cognitive ability, using the Montreal Cognitive Assessment (MoCA).

Overall, this work shows that information decomposition methods hold great promise for the development of explainable biomarkers of cognitive decline, and we expect that relating information decomposition and specific cognition will be a highly insightful research trajectory, with especially promising insights for cognitive disorders and neuronal disease.

## 5 Acknowledgements

The authors would like to thank Mar Estarellas, Tristán Bekinschtein, Andrés Canales-Johnson and the rest of the Cambridge Consciousness and Cognition Lab for their helpful input on this work and future directions. We would also like to thank Andrea Luppi and Silvia Rognone for their continued contributions on the trajectory of this work.

Data collection and sharing for the Alzheimer’s Disease Neuroimaging Initiative (ADNI) is funded by the National Institute on Aging (National Institutes of Health Grant U19 AG024904). The grantee organization is the Northern California Institute for Research and Education. In the past, ADNI has also received funding from the National Institute of Biomedical Imaging and Bioengineering, the Canadian Institutes of Health Research, and private sector contributions through the Foundation for the National Institutes of Health (FNIH) including generous contributions from the following: AbbVie, Alzheimer’s Association; Alzheimer’s Drug Discovery Foundation; Araclon Biotech; BioClinica, Inc.; Biogen; Bristol-Myers Squibb Company; CereSpir, Inc.; Cogstate; Eisai Inc.; Elan Pharmaceuticals, Inc.; Eli Lilly and Company; EuroImmun; F. Hoffmann-La Roche Ltd and its affiliated company Genentech, Inc.; Fujirebio; GE Healthcare; IXICO Ltd.; Janssen Alzheimer Immunotherapy Research & Development, LLC.; Johnson & Johnson Pharmaceutical Research & Development LLC.; Lumosity; Lundbeck; Merck & Co., Inc.; Meso Scale Diagnostics, LLC.; NeuroRx Research; Neurotrack Technologies; Novartis Pharmaceuticals Corporation; Pfizer Inc.; Piramal Imaging; Servier; Takeda Pharmaceutical Company; and Transition Therapeutics.

## Footnotes

1 For our analyses we have assumed distribution stationarity. As such, copy and erasure are linked; copy went up and not erasure, there would be a cascading imbalance in redundant representations, eventually corresponding to more information than the brain can physically represent. As such, erasure must also simultaneously increase. A similar effect links upwards and downwards causation.

