## Supplementary Materials for "Synergistic and redundant information dynamics are modulated by Alzheimer’s disease and cognitive impairment"

### Supplementary Methods

##### Anatomical Data Preprocessing (fMRIPrep boilerplate)

For each participant, all T1-weighted images were corrected for intensity non-uniformity (INU) with **N4BiasFieldCorrection** [63], distributed with ANTs 2.3.3 [1]. The T1w-reference was then skull-stripped with a Nipype implementation of the **antsBrainExtraction.sh** workflow (from ANTs), using OASIS30ANTs as the target template. Brain tissue segmentation of the cerebrospinal fluid (CSF), white-matter (WM) and gray-matter (GM) was performed on the brain-extracted T1-weighted image using **fast** [73]. An anatomical T1w-reference map was computed after registration the T1-weighted images (after INU-correction) using **mri_robust_template** [52]. Brain surfaces were reconstructed using **recon-all** [12], and the previously estimated brain mask was refined with an fMRIPrep custom variation of the method to reconcile ANTs-derived and FreeSurfer-derived segmentations of the cortical gray-matter of Mindboggle [33]. Volume-based spatial normalization to two standard spaces (MNI152Lin, MNI152NLin2009cAsym) was performed through nonlinear registration with **antsRegistration** (ANTs 2.3.3), using brain-extracted versions of both T1w reference and the T1w template. The following templates were were selected for spatial normalization and accessed with *TemplateFlow* 23.0.0 [10]: **Linear ICBM Average Brain (ICBM152) Stereotaxic Registration Model** [[42]; TemplateFlow ID: MNI152Lin], **ICBM 152 Nonlinear Asymmetrical template version 2009c** [[19]; TemplateFlow ID: MNI152NLin2009cAsym].

##### Functional data preprocessing (fMRIPrep boilerplate)

Preprocessing was performed across all BOLD runs from all participants across all sessions. First, a reference volume and its skull-stripped version were generated using a custom method from **fMRIPrep**. Head-motion parameters were estimated with respect to the BOLD reference (transformation matrices, and six corresponding rotation and translation parameters) before any spatiotemporal filtering using **mcflirt** [32]. BOLD runs were slice-time corrected where slice order was available from header data using **3dTshift** from AFNI [11]. The BOLD time-series (including slice-timing correction when applied) were resampled onto their original, native space by applying the transforms to correct for head-motion. We refer to these resampled BOLD time-series as *preprocessed BOLD in original space*, or just *preprocessed BOLD*. The BOLD reference was then co-registered to the T1w reference using **bbregister** (FreeSurfer) which implements boundary-based registration [26]. Co-registration was configured with six degrees of freedom. Several confounding time-series were calculated based on the *preprocessed BOLD*: framewise displacement (FD), DVARS, and three region-wise global signals. FD was computed using two formulations following Power (absolute sum of relative motions, [50]) and Jenkinson (relative root mean square displacement between affines, [32]). FD and DVARS were calculated for each functional run, both using their implementations in **Nipype** [following the definitions by 50]. The three global signals were extracted within the CSF, the WM, and the whole-brain masks. Additionally, a set of physiological regressors were extracted to allow for component-based noise correction [*CompCor*, 4]. Principal components were estimated after high-pass filtering the *preprocessed BOLD* time-series (using a discrete cosine filter with 128s cut-off) for the two *CompCor* variants: temporal (tCompCor) and anatomical (aCompCor). tCompCor components are then calculated from the top 2% variable voxels within the brain mask. For aCompCor, three probabilistic masks (CSF, WM and combined CSF+WM) are generated in anatomical space. The implementation differs from that of Behzadi et al. in that instead of eroding the masks by 2 pixels on BOLD space, a mask of pixels that likely contain a volume fraction of GM is subtracted from the aCompCor masks. This mask is obtained by dilating a GM mask extracted from the FreeSurfer's **aseg** segmentation, and it ensures components are not extracted from voxels containing a minimal fraction of GM. Finally, these masks are resampled into BOLD space and binarized by thresholding at 0.99 (as in the original implementation). Components are also calculated separately within the WM and CSF masks. For each CompCor decomposition, the $k$ components with the largest singular values are retained, such that the retained components' time series are sufficient to explain 50 percent of variance across the nuisance mask (CSF, WM, combined, or temporal). The remaining components are dropped from consideration. The head-motion estimates calculated in the correction step were also placed within the corresponding confounds file. The confound time series derived from head motion estimates and global signals were expanded with the inclusion of temporal derivatives and quadratic terms for each [58]. Frames that exceeded a threshold of 0.5 mm FD or 1.5 standardized DVARS were annotated as motion outliers. Additional nuisance timeseries are calculated by means of principal components analysis of the signal found within a thin band (*crown*) of voxels around the edge of the brain, as proposed by [49]. The BOLD time-series were resampled into standard space, generating a *preprocessed BOLD run in MNI152Lin space*. First, a reference volume and its skull-stripped version were generated using a custom methodology of **fMRIPrep**. All resamplings can be performed with *a single interpolation step* by composing all the pertinent transformations (i.e. head-motion transform matrices, susceptibility distortion correction when available, and co-registrations to anatomical and output spaces). Gridded (volumetric) resamplings were then performed using **antsApplyTransforms** (ANTs), configured with Lanczos interpolation to minimize the smoothing effects of other kernels [35]. Non-gridded (surface) resamplings were performed using **mri_vol2surf** (FreeSurfer).

#### Grouping into Montreal Cognitive Assessment categories

For all results presented in the main body of this paper utilising diagnostic categories (CN, MCI, AD), we performed similar analyses by grouping by Montreal cognitive assessment scores [47]. These groups were High MoCA: 29-30, Medium MoCA: 20-28, and Low MoCA: 0-19. These thresholds were chosen so as to approximately capture 15%, 70% and 15% of the dataset respectively. The total number of scans within each category were as follows:

| | High MoCA | Medium MoCA | Low MoCA |
|---|---|---|---|
| Participants | 131 | 775 | 181 |

#### Regional relative synergy and redundancy plots

In result 6.3 we plotted the absolute summed measures Synergy($R^i$) and Redundancy($R^i$) and subtracted each region's three-group mean so that increases in redundancy and decreases in synergy are easier to compare. We then ordered regions based on the difference between the change in the absolute synergy (redundancy) between the CN and AD groups to again highlight rough alignment with a principal unimodal-transmodal axis.

Due to computational constraints, significant differences were estimated at the $\alpha = 0.05$ level in a two-tailed $t$-test rather than using a permutation test. This was adjusted using an FDR correction. Significant differences were placed in chromosome plots underneath the main plot, constructed in python with Seaborn and Matplotlib.

##### Intranetwork atomic average calculations

For result 3.3 and 6.8, instead of summing across all pairs of regions, we took an average for each atom across all intranetwork interactions. To do this, we masked each of the matrices computed so as to obtain square subsets corresponding to interactions between pairs of ROIs inside of each Yeo network. These values were used as inputs to the taxonomic calculation for the result 3.3.

In the additional result 6.8, we applied independent $t$-tests with an FDR correction to test for significant differences between the cognitively normal (CN) and AD group. Results reported in 6.8 were transformed into Cohen's $d$ scores to allow for a normalised comparison of effect size. Only results surviving the significance threshold $\alpha = 0.05$ are reported in figure 12.

##### Synergy-minus-redundancy rank gradient

For supplementary result 6.10, we follow Luppi et al. [39] in computing a synergy-minus-redundancy rank gradient (seen in figure 15). Firstly, we found the regional summed redundancy (Redundancy($R^i$)) and summed synergy (Synergy($R^i$)) values, before ranking each region based on their synergy and their redundancy. Regions with the greatest amount of information in a given atom across all brain regions would be ranked the highest and vice versa.

We then placed each region in the synergy-redundancy hierarchy by subtracting the redundancy rank from the synergy rank:

$$w(R^i) = \text{rank}(\text{Synergy}(R^i)) - \text{rank}(\text{Redundancy}(R^i)).$$

The rank gradient $w(R^i)$ is positive when a region is highly synergistic but not offset by being highly redundant, and vice versa. Regions were plotted according to where they fell in the CN synergy-redundancy rank gradient.

To explore how this hierarchy might be destablised when comparing CN to AD, we computed Pearson correlations between the synergy-redundancy rank gradients between groups. To construct confidence intervals we performed stratified bootstrap resampling with $n = 1,000$ resamples (due to computational constraints more resampling was not feasible).

### 6 Additional Results

#### 6.1 Regional and network variation in summed information dynamics across cognitive score categories

Unless otherwise stated, all $p$-values in this section have been subject to an FDR correction via a Benjamini-Hochberg procedure.

##### 6.1.1 Heterogeneity in changes to ROI information dynamics between high and low MoCA

We applied an FDR-corrected Welch's $t$-test to compare the high and low MoCA categories, where we found significant decreases at the $p < 0.05$ level in synergy in all 30 ROIs in the ECN, 21/22 regions from the salience network, 40/46 regions in the default mode network, 27/32 regions in the subcortex, 21/26 regions in the dorsal attention network, 27/35 regions in the somatomotor network, 22/29 regions in the visual system, and 6/12 in the limbic system.

Similarly, when comparing redundancy in the high and low MoCA categories, significant increases at the $p < 0.05$ level were found in 29/30 ROIs in the ECN, 24/32

ROIs in the subcortex, 32/46 ROIs in the DMN, 15/26 ROIs in the dorsal attention network, 11/22 ROIs in the salience network, 5/12 ROIs in the limbic system, 10/29 ROIs in the visual system, and 4/35 ROIs in the somatomotor network.

No comparisons were significant for comparisons between the high and medium MoCA group after applying the FDR correction.

###### 6.1.2 Organisation of changes in summed information dynamics across cognitive scores

To compare against a cognitive score, we also grouped participants into three categories based on their MoCA score (combining data from across all diagnostic groups). Highest-scoring participants (MoCA 29-30) formed our normal cognition category. Medium-scoring participants (MoCA 20-28) formed our medium cognition category. Low-scoring participants (MoCA 0-19) formed our low cognition category. We then performed the same $z$-scoring procedure as with the diagnostic groups before computing the first principal component of variation as before. The results are shown in figure 4.

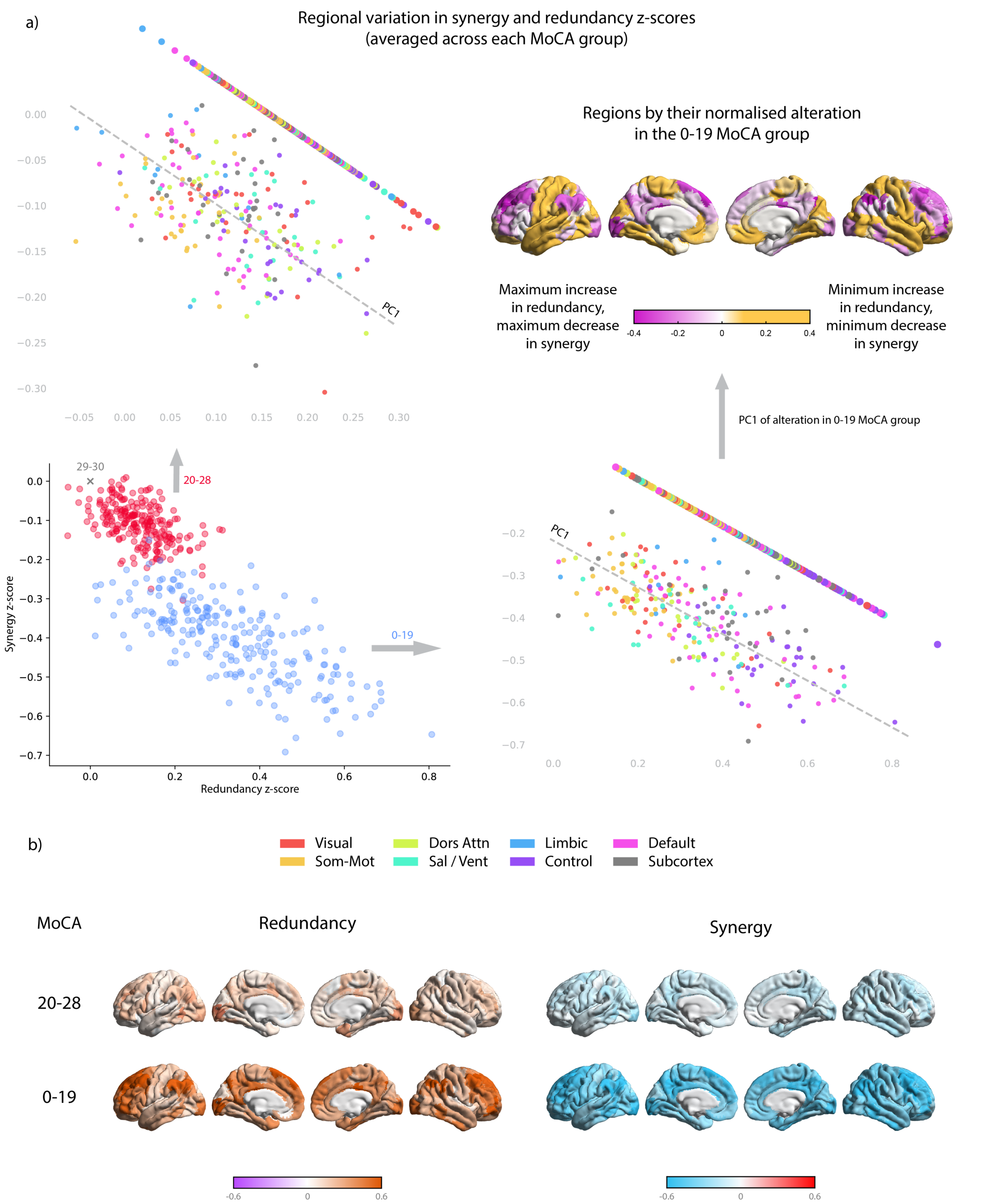


**Figure 4. Information dynamics change in correspondence with cognitive decline**. As per figure 1, but using Montreal Cognitive Assessment scores (Low MoCA: 0-19, Medium MoCA: 20-28, High MoCA: 29-30). In a), the average regional changes in information dynamics for each cognitive group were $z$-scored based on their mean in the high MoCA (cognitively highest-functioning) group (MoCA 29-30). These were then projected onto their first principal component to illustrate the relationship between redundancy and synergy change across brain regions. In figure b) the $z$-scores of redundancy and synergy are shown in comparison to the MoCA 29-30 group.

Results are similar in fashion to the diagnostic categories but with a subtle difference for the medium cognition group (MoCA 20-28). The amount of variance explained by the first principal component decreased when using cognitive scores compared to the diagnostic label, with PC1 explaining 75.6% of the variance in the medium MoCA group and 88.6% of the variance in the low MoCA group.

#### 6.2 Tables of network $z$-scores comparing CN to MCI, AD and high MoCA to medium, low MoCA

| Group | Vis | SomMat | DA | Sal | Limbic | ECN | DMN | Subctx | Global |
|---|---|---|---|---|---|---|---|---|---|
| MCI | 0.01 | -0.08 | -0.02 | -0.05 | 0.10 | 0.07 | 0.04 | 0.09 | 0.017 |
| AD | 0.20 | 0.11 | 0.22 | 0.18 | 0.36 | 0.43 | 0.37 | 0.37 | 0.279 |

**Table 1.** Redundancy $z$-scores in the MCI and AD groups, averaged across 7 Yeo networks and additional subcortical regions.

| Group | Vis | SomMat | DA | Sal | Limbic | ECN | DMN | Subctx | Global |
|---|---|---|---|---|---|---|---|---|---|
| MCI | -0.10 | -0.05 | -0.09 | -0.06 | -0.13 | -0.13 | -0.12 | -0.13 | -0.102 |
| AD | -0.44 | -0.37 | -0.43 | -0.42 | -0.44 | -0.54 | -0.49 | -0.48 | -0.456 |

**Table 2.** Synergy $z$-scores in the MCI and AD groups, averaged across 7 Yeo networks and additional subcortical regions.

| Group | Vis | SM | DA | Sal | Limb | ECN | DMN | Subctx | Gbl |
|---|---|---|---|---|---|---|---|---|---|
| MoCA 20-28 | 0.14 | 0.07 | 0.13 | 0.15 | 0.07 | 0.15 | 0.10 | 0.11 | 0.115 |
| MoCA 0-19 | 0.29 | 0.18 | 0.29 | 0.31 | 0.31 | 0.50 | 0.37 | 0.39 | 0.334 |

**Table 3.** Redundancy $z$-scores, averaged across 7 Yeo networks and additional subcortical regions across both MoCA categories as compared to the high MoCA category.

| Group | Vis | SM | DA | Sal | Limb | ECN | DMN | Subctx | Gbl |
|---|---|---|---|---|---|---|---|---|---|
| MoCA 20-28 | -0.10 | -0.10 | -0.12 | -0.13 | -0.07 | -0.13 | -0.10 | -0.10 | -0.109 |
| MoCA 0-19 | -0.37 | -0.34 | -0.39 | -0.41 | -0.34 | -0.50 | -0.43 | -0.40 | -0.402 |

**Table 4.** Synergy $z$-scores, averaged across 7 Yeo networks and additional subcortical regions across both MoCA categories as compared to the high MoCA category.

#### 6.3 Further illustrations demonstrating absolute changes in synergy and redundancy

To further expose these differences, we looked not only at $z$-scored regional changes, but also changes in total synergy and redundancy (in bits) between regions across diagnostic groups and cognitive score categories. For each region, the average total synergy and redundancy with all other regions was computed, and the brain regions ordered based on the difference between the total in the CN and dementia groups (AD), as shown in figure 5.

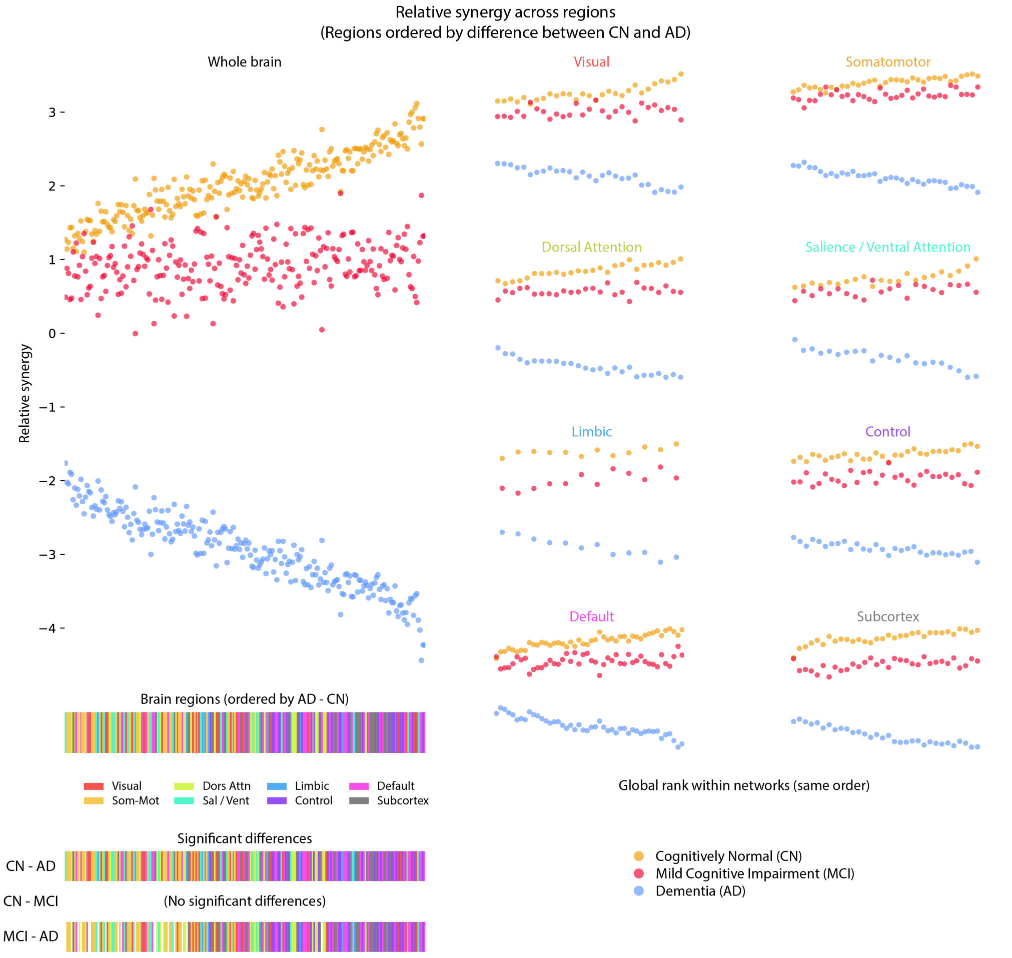


**Figure 5. Multimodal regions in the CN brain lose the most total synergy when compared to individuals with MCI or Alzheimer's disease (AD)**. Left: all brain regions ordered by their synergistic decline from CN to AD, subtracting the mean synergy for that region across all three groups for readability. Bottom left: significant differences between groups in each region $t$-tested and FDR corrected at the $\alpha = 0.05$ level. Their relative total synergy (in bits) is plotted against the three-group mean. Right: the same result, but separated out into the 7 Yeo networks (plus subcortex).

In regions which have the most unimodal processing, such as those in visual or somatomotor regions, the decrease in synergy appears visibly smallest in the plot, while the synergy decrease is more visible in the control and default mode networks, along with subcortical regions, confirming earlier statistical tests. After correcting for multiple comparisons with an FDR correction, we found no statistically significant differences between synergy in the CN and MCI group at the $\alpha = 0.05$ level. Between the CN and AD group we found *all* regions (232) met the threshold for significance. Between the MCI and AD group, 213 regions met the threshold for significance, as shown in the bottom left of figure 5.

In order to confirm that this is also modulated by cognition, we also applied this to our three MoCA groups (29-30: High, 20-28: Medium, and 0-19: Low). In figure 6 we plot regions based on the difference in synergistic information between the high and low MoCA groups.

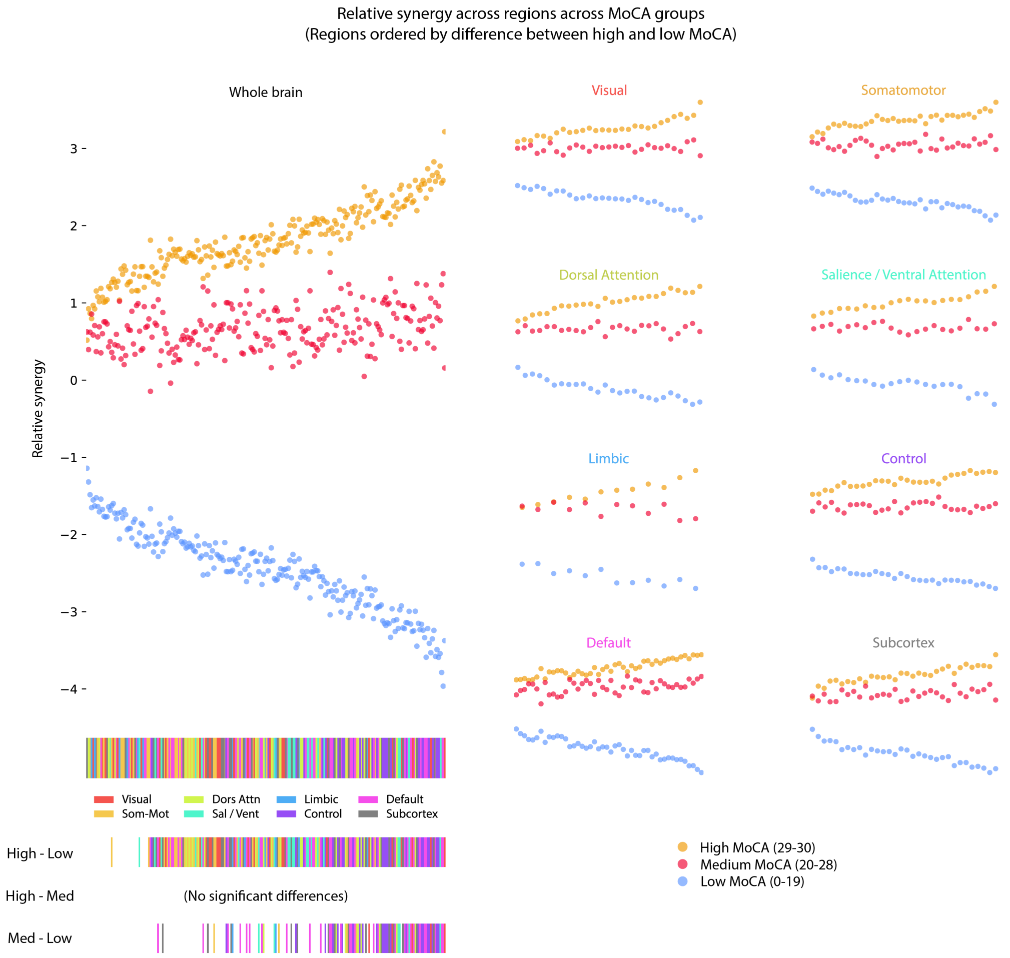


**Figure 6. MoCA scores separate regions by relative synergy**. Left: all brain regions ordered by the difference in synergy between the high and low MoCA groups, subtracting the mean synergy for that region across all three groups for readability. Bottom left: significant differences between groups in each region $t$-tested and FDR corrected at the $\alpha = 0.05$ level. Right: subsets of the plot depicting the 7 Yeo networks (plus subcortex).

In addition to synergy dynamics, we also investigated redundancy dynamics across regions, which we show in figure 7. In contrast to the diagnostic groups, where MCI demonstrated substantial overlap with the CN group, we saw that our intermediate cognition group (red, MoCA 20-28) exhibited redundancy between those of the low (blue) and high (yellow) cognition groups.

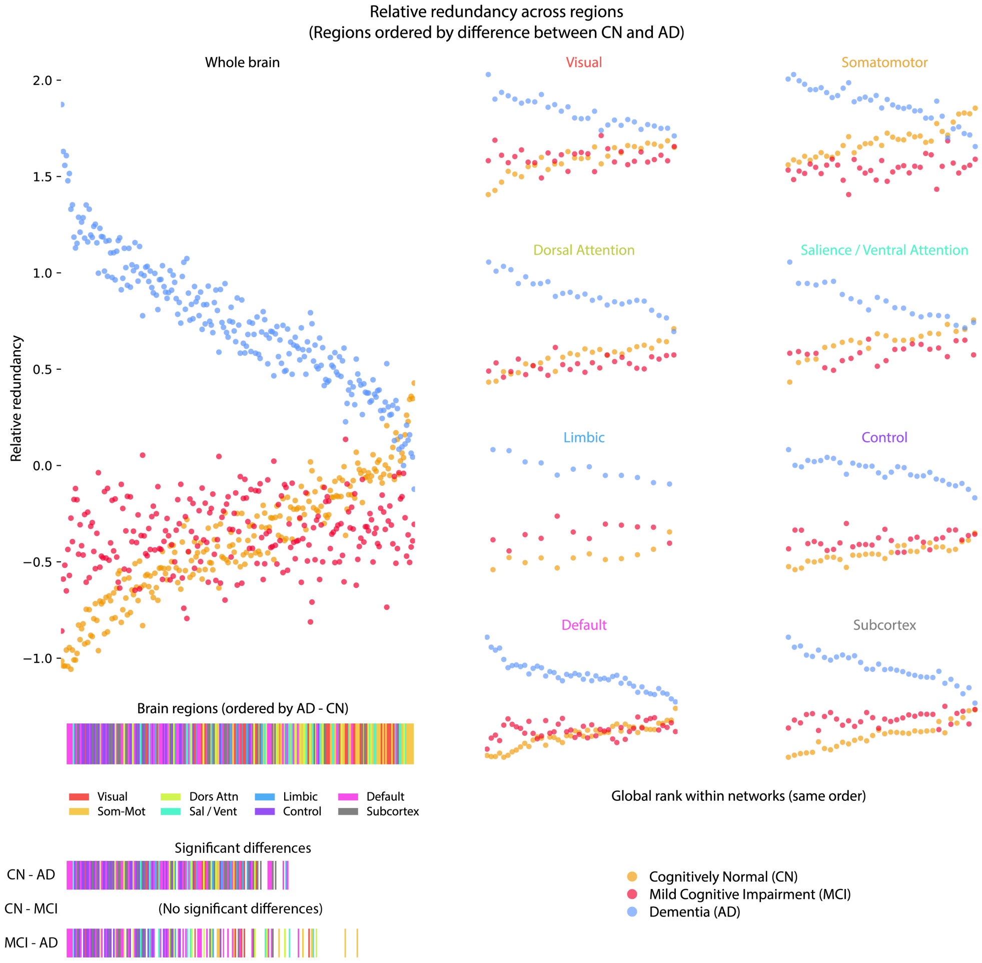


**Figure 7. Redundant information changes are non-homogeneous in MCI, but global in AD** Top left: as with the previous figure, the total redundancy is plotted for each region with that region's mean across the three groups subtracted for ease of readability. Regions are ordered according to the size of the difference between the CN and AD group. Bottom left: significant differences $t$-tested at the $\alpha = 0.05$ level using FDR correction. Right: regions with the same order as they appear in the whole brain, but segregated into different Yeo networks (along with subcortical regions).

In contrast to the altered synergistic signature in the average AD brain, which decreased when compared to CN in all regions (with this pattern also holding in MCI), the redundant information has slightly more complex behaviour. In just under half of brain regions (predominantly unimodal, somatosensory regions), redundancy saw a decrease in MCI compared to the CN group, yet saw an increase in the Alzheimer's disease (AD) group when compared to CN, possibly highlighting a compensatory information processing mechanism. The greatest increase from CN to AD, meanwhile, was observed in the control and default mode networks, though this distinction remains unclear in the MCI, where the clearest changes are in limbic and subcortical areas.

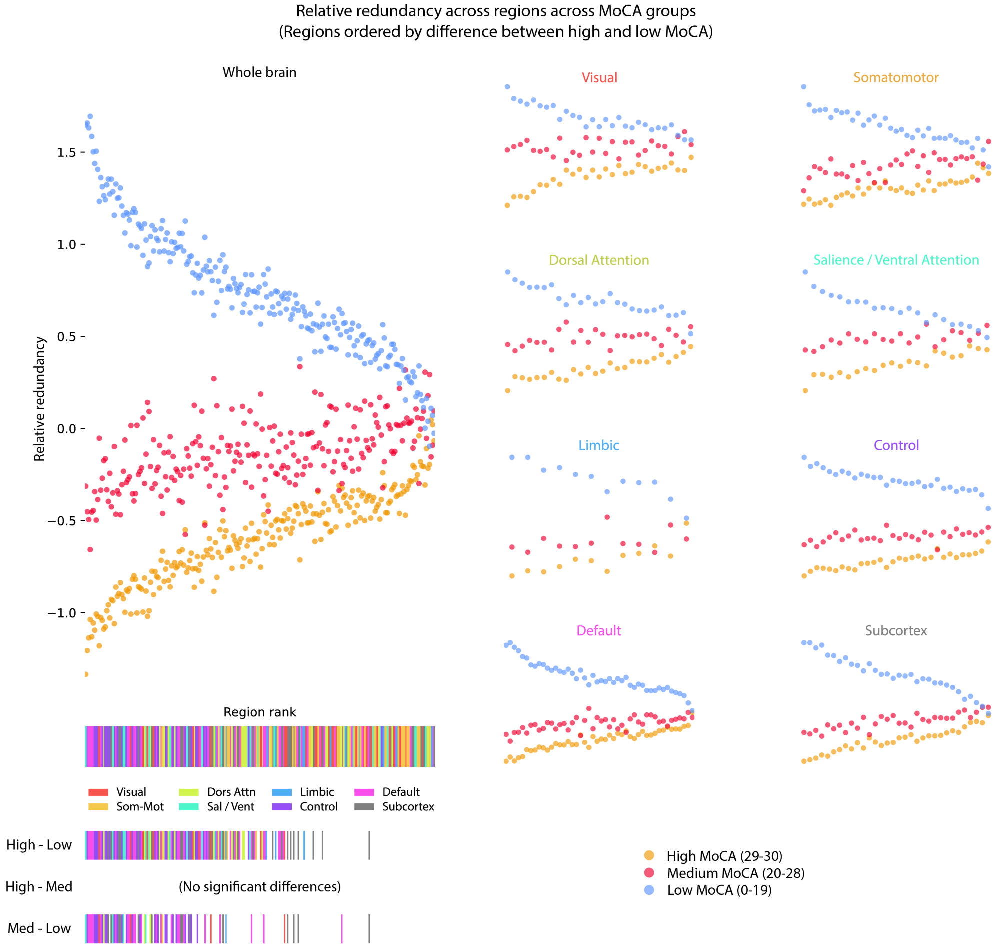


**Figure 8. Redundant interactions increased with cognitive decline** Top left: as with the previous figure, the total redundancy is plotted for each region with that region's mean across the three groups subtracted for readability. Regions are ordered according to the size of the difference between the high MoCA and low MoCA groups. Bottom left: significant differences $t$-tested at the $\alpha = 0.05$ level using FDR correction. Right: regions with their same location as in the whole brain, but separated into different Yeo networks (along with subcortical regions).

#### 6.4 Synergy loss and redundancy gain in low cognition are not uniform across networks

##### 6.4.1 Network-level differences in summed synergy and redundancy between High and Medium MoCA

None of the ROIs exhibited a significant difference in either synergy or redundancy between the high and medium MoCA categories at the 0.05 significance level. Similarly none of the networks exhibited significant differences in either synergy or redundancy between high and medium MoCA at the 0.05 significance level.

##### 6.4.2 Network-level differences between summed synergy and redundancy between High and Low MoCA

When taking averages of the total summed synergy across each network, we found significant decreases in synergy across all networks when comparing the High MoCA (29-30) and Low MoCA (0-19) groups at the 0.05 level. Effect sizes were largest in the control (Welch's $t = -3.472$, FDR-adjusted $p = 0.005$, Hedge's $g = -0.363$), default mode ($t = -3.033$, $p = 0.007$, $g = -0.318$), salience ($t = -2.928$, $p = 0.007$, $g = -0.308$), and subcortical regions ($t = -2.928$, $p = 0.007$, $g = -0.307$). Smaller but still significant effects were found in the dorsal attention ($t = -2.764$, $p = 0.010$, $g = -0.292$), visual ($t = -2.686$, $p = 0.010$, $g = -0.284$), somatomotor ($t = -2.506$, $p = 0.014$, $g = -0.265$), and limbic networks ($t = -2.476$, $p = 0.014$, $g = -0.265$).

When looking at changes in total summed redundancy between the High MoCA and Low MoCA groups we found significant increases in the control (Welch's $t = 4.424$, FDR-adjusted $p < 0.001$, Hedges' $g = 0.477$), subcortical ($t = 3.496$, $p = 0.002$, $g = 0.377$), default mode ($t = 3.347$, $p = 0.002$, $g = 0.363$), and salience networks ($t = 2.861$, $p = 0.009$, $g = 0.315$). Smaller but still significant increases in redundancy were also found in the dorsal attention ($t = 2.661$, $p = 0.013$, $g = 0.297$), visual ($t = 2.606$, $p = 0.013$, $g = 0.288$), and limbic networks ($t = 2.534$, $p = 0.013$, $g = 0.0267$). The somatomotor regions did not exhibit a statistically significant increase in redundancy.

##### 6.4.3 Changes in information dynamics corresponding to cognitive score are not solely explained by a global effect

In result 3.2.1, we verified that changes in network information dynamics could not be explained by a global effect alone. We verified whether this hypothesis also holds across MoCA score categories.

When comparing synergy dynamics in the low MoCA group (0-19) against the high MoCA group (29-30) with the summed synergy measure, we found that the executive control network showed a greater decrease in synergy than could be explained by a global effect ($\Delta\text{syn} = -5.121$, $p < 0.001$), while the somatomotor ($\Delta\text{syn} = -3.745$, $p = 0.001$) and limbic systems ($\Delta\text{syn} = -3.655$, $p = 0.045$), while showing a decrease in synergy, exhibited less of a decrease than could be explained by a global effect.

When comparing redundancy dynamics in the same two groups, we found that only the ECN ($\Delta\text{red} = 1.941$, $p < 0.001$) showed an increase greater than a global effect would allow, and the somatomotor network ($\Delta\text{red} = 0.937$, $p < 0.001$) showed an increase smaller than a global effect would explain. Significant deviations from a global effect are shown in figure 9.

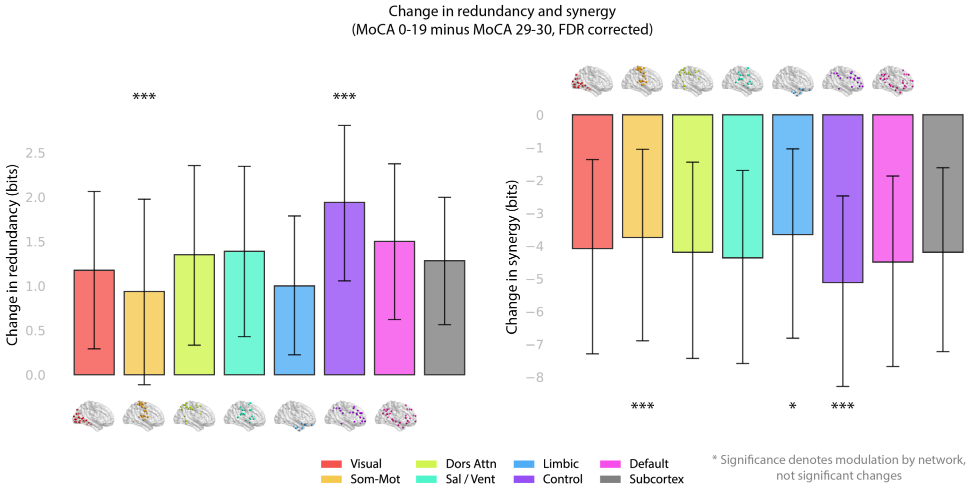


**Figure 9. Somatomotor and control regions exhibit specific changes in redundancy and synergy when compared between high and low MoCA groups.** As with figure 2, the total redundancy and synergy is added up for each region before being averaged across each network to find the average total changes in that network's regional redundancy and synergy when comparing high and low MoCA categories. Left: increases in total redundancy averaged across each of the Yeo networks (plus subcortex); right: decreases in total synergy averaged across each of the Yeo networks. As with figure 2, significance was found using a permutation test with $n = 10,000$ resamples and FDR correction to account for multiple comparisons, shuffling network labels to check for inhomogeneous information dynamical changes with a change in cognition. Significances are illustrated at the $\alpha = 0.05^{*}, 0.01^{**}$, and $0.001^{***}$ levels respectively. Confidence intervals were computed by bootstrap resampling to obtain the 95% confidence interval.

#### 6.5 Atomic modes vary across MoCA groups

For all significance tests in this section we applied the Benjamini-Hochberg procedure for false discovery rate correction.

Because we observed differences in the information theoretic profiles of the MoCA groups in result 3.1, we also applied the mode analysis to MoCA classes. As with the diagnostic categories, we applied a one-way FDR-corrected ANOVA across the three cognitive categories (High MoCA, Medium MoCA, and Low MoCA). The ANOVA flagged a group-wise variation in the multi-scale causation mode in the ECN ($p = 0.001$), DMN ($p = 0.019$), and subcortex ($p = 0.008$), along with changes in the copy-erasure mode in the ECN ($p = 0.008$) and subcortex ($p = 0.037$). The storage mode also saw some significant variation in the ECN ($p = 0.001$), DMN ($p = 0.008$), and the subcortex ($p = 0.008$), with the transfer atom only showing group variation in the ECN ($p = 0.034$). Notably the ECN was the only network to consistently reach the significance threshold across all modes. Results are shown in figure 10.

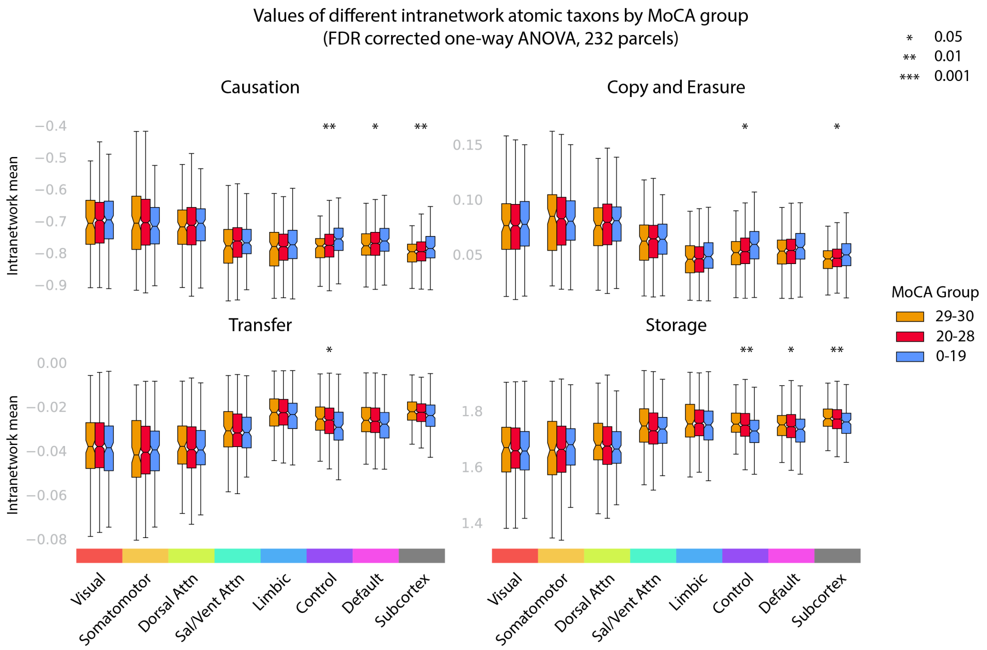


**Figure 10. Qualitative information processing is modulated by MoCA score group.** As with figure 3, the taxonomic information processing was computed for all pairs of regions within each of the 7 Yeo networks (along with additional subcortical regions). For each combination of network and four ΦID-modes (where we have collapsed upwards and downwards causation into one mode and copy and erasure into one mode), a one-way ANOVA was performed with a Benjamini-Hochberg correction for the false discovery rate.

To work out what drove these group variations, we performed a post-hoc Welch $t$-test to the MoCA categories, with results shown in table 6. No significant differences were identified between the high and medium cognition groups. The largest effect (in all modes) was observed in the ECN, with the greatest decrease observed in the storage mode.

#### 6.6 Post-hoc Welch's $t$-tests for modes (diagnosis and MoCA)

| Mode | Network | Comparison | Mean diff | Welch $t$ | FDR Adj $p$ | Sig |
|---|---|---|---|---|---|---|
| Causation | Control | CN vs AD | 0.041 | 4.037 | $< 0.001$ | *** |
| Causation | Control | MCI vs AD | 0.034 | 3.295 | 0.002 | ** |
| Causation | Default | CN vs AD | 0.030 | 3.047 | 0.008 | ** |
| Causation | Default | MCI vs AD | 0.027 | 2.695 | 0.011 | * |
| Causation | Limbic | CN vs AD | 0.033 | 3.168 | 0.005 | ** |
| Causation | Limbic | MCI vs AD | 0.025 | 2.347 | 0.030 | * |
| Causation | SubCtx | CN vs AD | 0.035 | 3.694 | 0.001 | *** |
| Causation | SubCtx | MCI vs AD | 0.027 | 2.722 | 0.010 | * |
| Causation | Visual | CN vs AD | 0.029 | 2.448 | 0.045 | * |
| Copy Erasure | Control | CN vs AD | 0.009 | 4.662 | $< 0.001$ | *** |
| Copy Erasure | Control | MCI vs AD | 0.007 | 3.722 | $< 0.001$ | *** |
| Copy Erasure | Default | CN vs AD | 0.006 | 3.181 | 0.005 | ** |
| Copy Erasure | Default | MCI vs AD | 0.006 | 2.961 | 0.005 | ** |
| Copy Erasure | Limbic | CN vs AD | 0.007 | 3.858 | $< 0.001$ | *** |
| Copy Erasure | Limbic | MCI vs AD | 0.005 | 2.749 | 0.009 | ** |
| Copy Erasure | SubCtx | CN vs AD | 0.008 | 4.799 | $< 0.001$ | *** |
| Copy Erasure | SubCtx | MCI vs AD | 0.005 | 3.025 | 0.004 | ** |
| Storage | Control | CN vs AD | -0.046 | -4.170 | $< 0.001$ | *** |
| Storage | Control | MCI vs AD | -0.038 | -3.404 | 0.001 | ** |
| Storage | Default | CN vs AD | -0.033 | -3.116 | 0.006 | ** |
| Storage | Default | MCI vs AD | -0.030 | -2.776 | 0.009 | ** |
| Storage | Limbic | CN vs AD | -0.036 | -3.276 | 0.004 | ** |
| Storage | Limbic | MCI vs AD | -0.028 | -2.431 | 0.024 | * |
| Storage | SubCtx | CN vs AD | -0.040 | -3.686 | 0.001 | *** |
| Storage | SubCtx | MCI vs AD | -0.030 | -2.709 | 0.011 | * |
| Storage | Visual | CN vs AD | -0.032 | -2.522 | 0.037 | * |
| Transfer | Control | CN vs AD | -0.004 | -4.274 | $< 0.001$ | *** |
| Transfer | Control | MCI vs AD | -0.003 | -3.376 | 0.001 | ** |
| Transfer | Default | CN vs AD | -0.003 | -2.775 | 0.016 | * |
| Transfer | Default | MCI vs AD | -0.002 | -2.577 | 0.016 | * |
| Transfer | Limbic | CN vs AD | -0.003 | -3.332 | 0.003 | ** |
| Transfer | Limbic | MCI vs AD | -0.002 | -2.304 | 0.033 | * |
| Transfer | SubCtx | CN vs AD | -0.003 | -2.986 | 0.009 | ** |

**Table 5.** Results of a post-hoc $t$-test to compare between groups after the initial one-way FDR adjusted ANOVA. Significance was also adjusted with an FDR correction. Only significant results are highlighted here.

| Mode | Network | Comparison | Mean diff | Welch $t$ | FDR Adj $p$ | Sig |
|---|---|---|---|---|---|---|
| Causation | Control | High vs Low | 0.038 | 3.292 | 0.003 | ** |
| Causation | Control | Med vs Low | 0.028 | 2.749 | 0.010 | ** |
| Causation | Default | High vs Low | 0.026 | 2.303 | 0.045 | * |
| Causation | Default | Med vs Low | 0.021 | 2.185 | 0.045 | * |
| Causation | SubCtx | High vs Low | 0.030 | 2.721 | 0.021 | * |
| Causation | SubCtx | Med vs Low | 0.022 | 2.319 | 0.032 | * |
| Copy Erasure | Control | Med vs Low | 0.006 | 3.095 | 0.007 | ** |
| Copy Erasure | Control | High vs Low | 0.007 | 2.868 | 0.007 | ** |
| Copy Erasure | SubCtx | High vs Low | 0.006 | 2.876 | 0.013 | * |
| Copy Erasure | SubCtx | Med vs Low | 0.004 | 2.477 | 0.021 | * |
| Storage | Control | High vs Low | -0.041 | -3.320 | 0.003 | ** |
| Storage | Control | Med vs Low | -0.031 | -2.840 | 0.007 | ** |
| Storage | Default | High vs Low | -0.029 | -2.291 | 0.039 | * |
| Storage | Default | Med vs Low | -0.023 | -2.238 | 0.039 | * |
| Storage | SubCtx | High vs Low | -0.033 | -2.713 | 0.021 | * |
| Storage | SubCtx | Med vs Low | -0.025 | -2.301 | 0.034 | * |
| Transfer | Control | High vs Low | -0.003 | -2.634 | 0.013 | * |
| Transfer | Control | Med vs Low | -0.003 | -2.762 | 0.013 | * |

**Table 6.** As with table 5, a post-hoc $t$-test was performed to compare between MoCA groups after the initial one-way FDR adjusted ANOVA. Significance was also adjusted with an FDR correction. Tests which did not meet the threshold for significance are not reported for space.

#### 6.7 Distinct information processing profiles with disease progression

Diagnostic groups had distinct information processing profiles when examined under integrated information decomposition ($\Phi$ID), as can be seen from contrasting the synergistic and redundant connectivity matrices. Figure 11 shows the interactions between pairs of brain regions for the a) Redundancy and b) Synergy atoms across the three diagnostic groups after averaging across the whole of the group. Differences between the CN and AD diagnostic groups are reported at the $\alpha = 0.05$ level using a two-tailed $t$-test with FDR correction (due to computational constraints a permutation test was not performed, c.f. section 2).

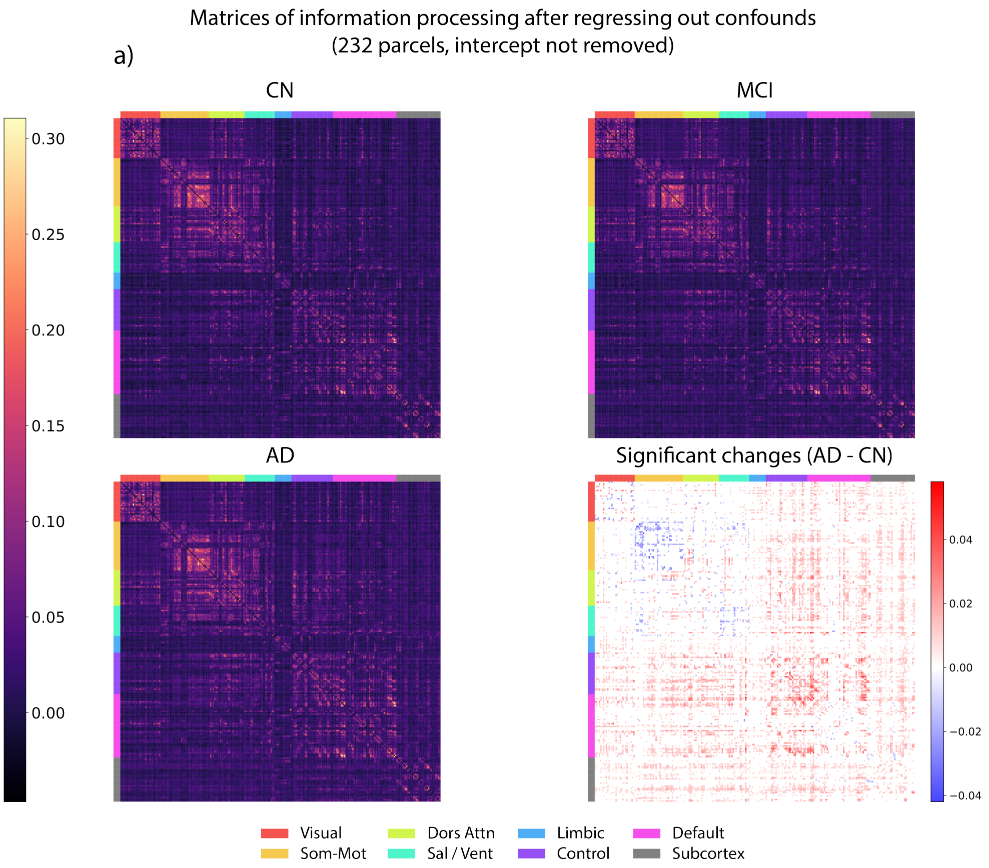

Matrices of information processing after regressing out confounds
(232 parcels, intercept not removed)
a)
CN
MCI
AD
Significant changes (AD - CN)
0.30
0.25
0.20
0.15
0.10
0.05
0.00
0.04
0.02
0.00
−0.02
−0.04
Visual
Som-Mot
Dors Attn
Sal / Vent
Limbic
Control
Default
Subcortex

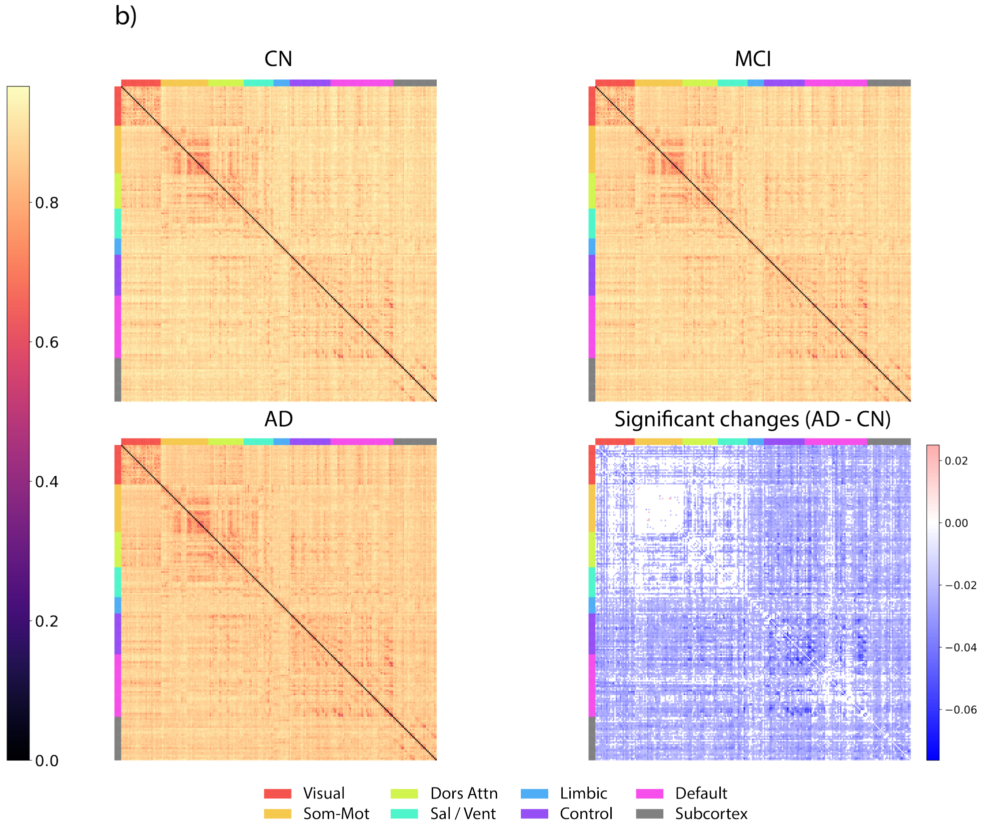


**Figure 11.** Matrices depicting the average **a)** Redundancy and **b)** Synergy ΦID atoms between each pair of 232 regions of interest (ROIs) taken from an extended Schaefer 200 atlas, re-ordered so that atoms in the same Yeo network are adjacent. Depicted is the residual value of the atom after regressing out age, gender, years of education and intracranial volume (ICV). Results are given across three diagnostic groups: CN, mild cognitive impairment (MCI) and AD. In addition to the atomic plots, in the bottom right we highlight significant differences in each atom between the CN and AD groups with a two-tailed $t$-test at the $\alpha = 0.05$ significance level, adjusting for the false discovery rate using the Benjamini-Hochberg procedure. Red areas denote significant increases in AD compared to the CN group, and blue areas denote significant decreases.

After applying the FDR correction, we found 39,988 of a possible 53,824 synergy atoms calculated exhibited a significant decrease in AD compared to CN at the $p < 0.05$ level using Welch's $t$-test. Only 10 pairs of ROIs saw interactions with a significant increase in synergy in AD compared to CN at the $p < 0.05$ level.

Similarly, 10,498 of a possible 53,824 pairs of redundancy atoms computed saw a statistically significant increase at the $p < 0.05$ level when using Welch's $t$-test with an FDR correction. Only 812 pairs saw a decrease in redundancy.

#### 6.8 Intranetwork representations track disease progression

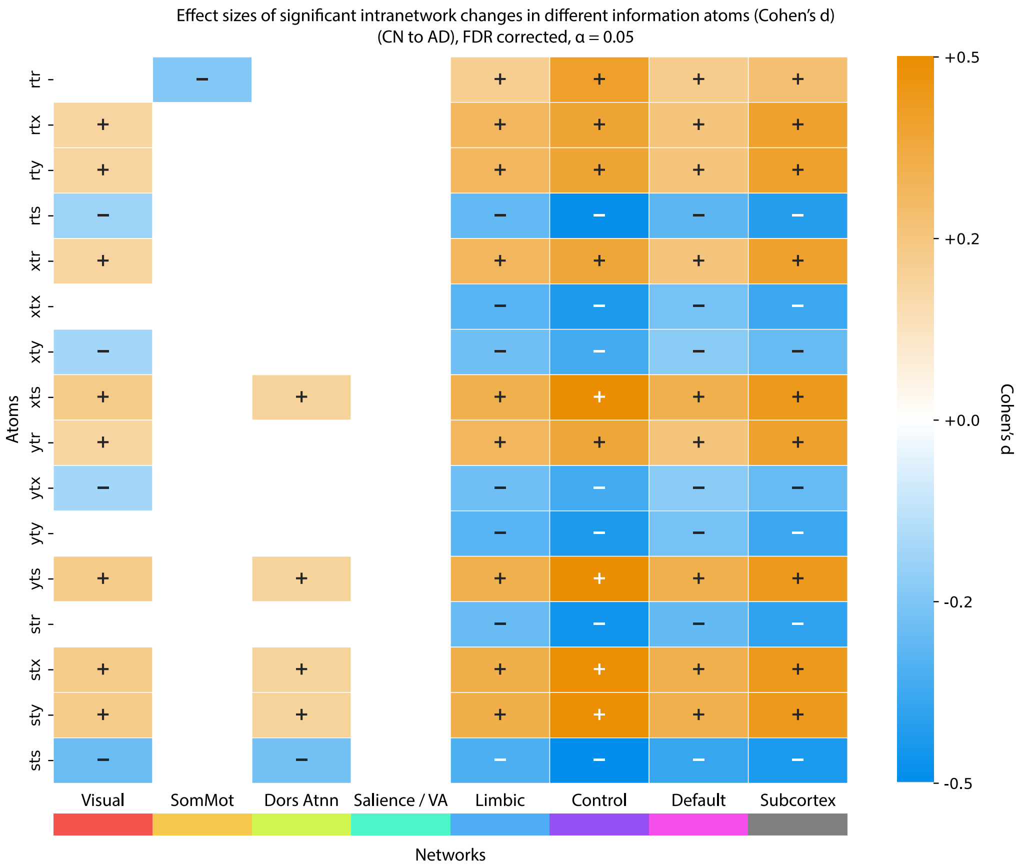


**Figure 12.** Significant intranetwork changes when comparing the CN group against those with AD. Each row corresponds to one of the 16 ΦID atoms, and each column to one of the Yeo networks (or subcortex). Interactions between all ROIs in a given Yeo network (or other subcortical regions) are averaged together to get a value for a network-wise intranetwork information processing. Changes are scaled using Cohen's $d$ as a normalised metric. Significant differences meeting the $\alpha = 0.05$ significance threshold using independent $t$-tests (with an FDR/Benjamini-Hochberg multiple comparisons correction) are depicted in orange for significant increases and blue for significant decreases.

As can be seen in figure 11 a), many of the atomic differences appear to be strongest when looking at intra-network differences, where the two ROIs being examined are taken from the same Yeo network [71]. Average ROI interactions for intra-network information dynamics are also altered in AD. Accounting for multiple comparisons with the Benjamini-Hochberg procedure across all 16 ΦID atoms and 8 macroscopic brain regions, we found that 82 network-atom pairs exhibited significant changes at the $\alpha = 0.05$ level in two-tailed $t$-tests out of a possible 128. Of these, 34 exhibited a significant *decrease* and 48 exhibited a significant *increase*.

Notably, with only one exception, atoms exhibited similar behaviour across all macroscopic brain regions, either increasing or decreasing in AD compared to CN. The only exception was the redundancy atom in the somatomotor network, which exhibited a decrease ($t = -2.838$, $p = 0.009$) despite showing an increase in the limbic ($t = 2.510$, $p = 0.022$), control ($t = 4.873$, $p < 0.001$), DMN ($t = 2.612$, $p = 0.017$), and subcortical areas ($t = 3.279$, $p = 0.003$).

Synergy saw significant intranetwork decreases in AD compared to CN in the visual ($t = -3.398$, $p = 0.002$), dorsal attention ($t = -3.149$, $p = 0.004$), limbic ($t = -4.092$, $p < 0.001$), control ($t = -5.842$, $p < 0.001$), default mode ($-4.540$, $p < 0.001$), and subcortical networks ($t = -5.236$, $p < 0.001$).

The macroscopic region with the strongest atomic changes overall was the ECN. The somatomotor and salience networks exhibited almost no changes at all when comparing the CN group against the AD group.

Importantly these changes were much more robust in the intranetwork version rather than the summed measure. That is to say, exploring intranetwork interactions showed much greater changes in redundancy and synergy than the summed measure in the main text.

#### 6.9 Alternative plots for result 6.3

When presenting additional result 6.3, we presented figures 5, 7, 6, and 8 by ordering all 232 regions according to the difference between the information quantity (synergy/redundancy) in AD minus CN. This was done after subtracting out the mean difference so that groups could more easily be contrasted. An alternative approach would be to order regions by their total synergy/redundancy in a particular group. Here we give the corresponding plots in figures 13 and 14. When looking at the total synergy and total redundancy, it is clear that the synergistic effects are much larger in absolute terms (ranging around 200 bits when summed across all regions, compared to about 5 bits in redundancy). As before, it can be seen that regions exhibit a global decrease in synergy.

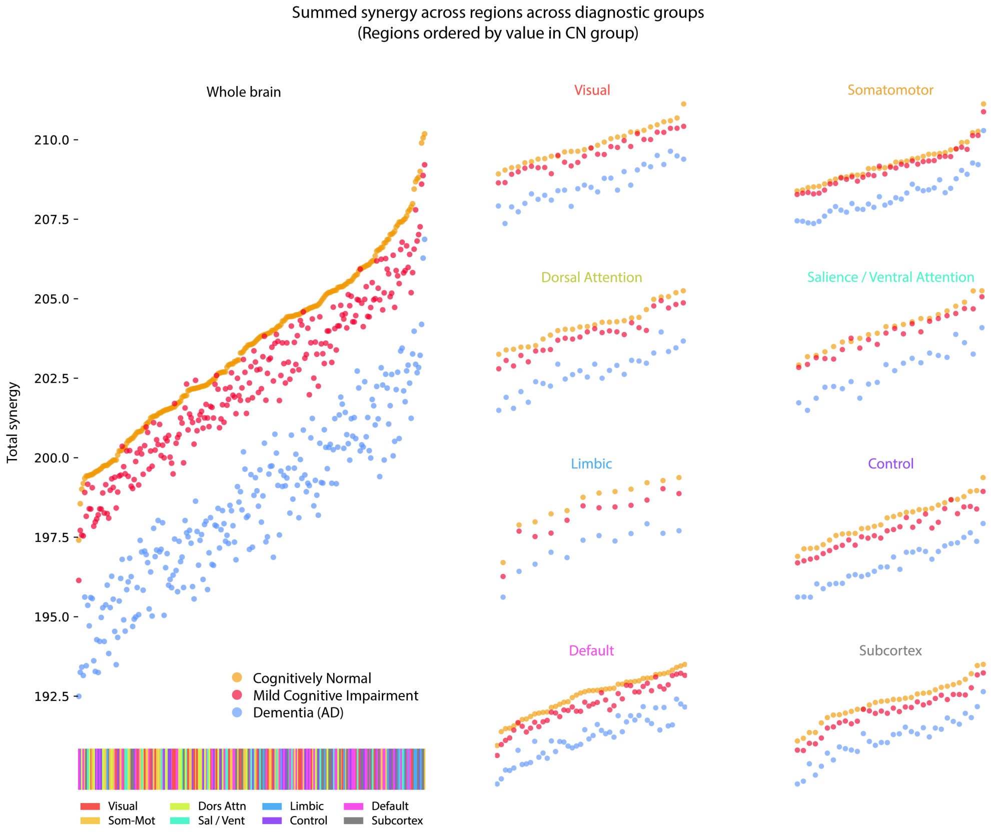

**Figure 13. Total synergy decreases with cognitive decline from CN to MCI to AD**. For each ROI in the Schaefer-232 atlas, that region's synergy with all other regions was summed to get a total synergy measure in bits. Left: all brain regions ordered by their total synergy in the CN group (orange), also showing that region's values in MCI (red) and AD (blue). Regions are coloured by their Yeo network. Right: the same regions restricted to each of the 7 Yeo networks (plus the subcortical regions).

We present the same plots here with the redundancy atom in figure 14. As expected due to result 3.1, there is significant overlap between the CN regions and the MCI regions. Notably, by looking at the subsets of regions on the right, it can be seen that the average group redundancy is slightly more separable between CN and MCI in the limbic system, ECN, and subcortex.

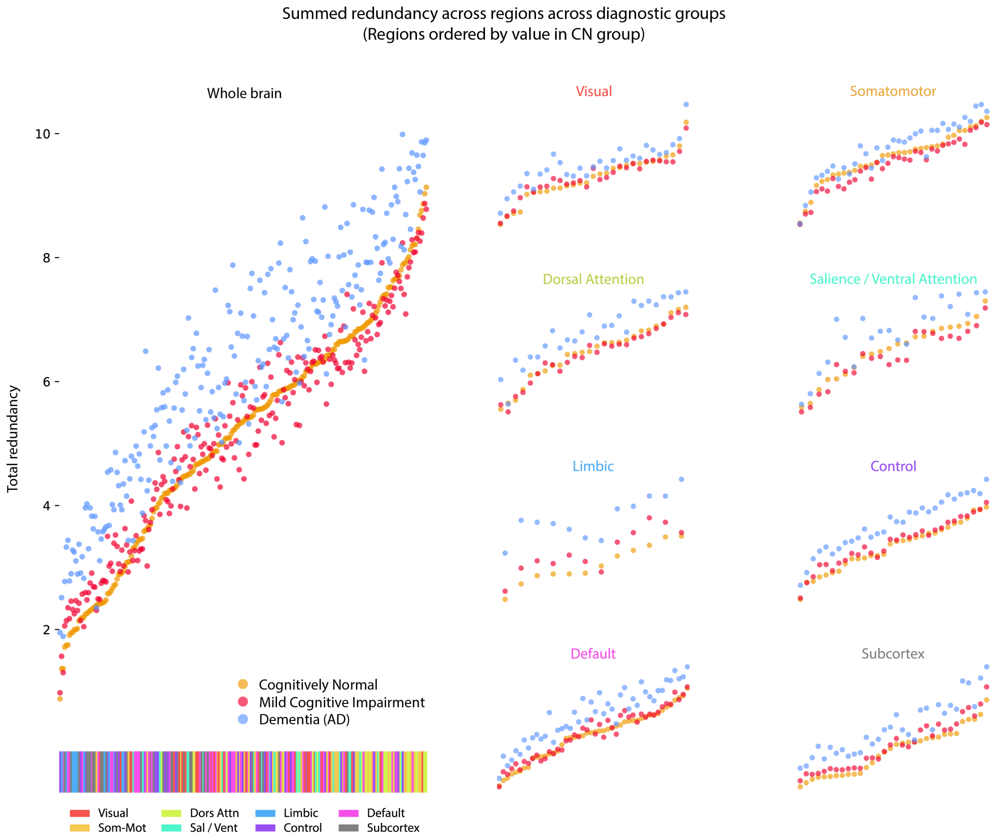


**Figure 14. Total redundancy increases in AD compared to CN, but MCI is difficult to separate.** As with figure 13, but with redundancy. Each region's total redundancy with all other regions was calculated and plotted in order of how much redundancy that region had in the CN group. Regions are coloured by Yeo network. Left: all brain regions ordered by CN. Right: brain regions from each of the 7 Yeo networks and subcortex, in the same order.

#### 6.10 Weak restructuring of the information processing hierarchy in AD

Following the information processing hierarchy described by Luppi et al. in 2022 [39], we ranked regions based on their temporal redundancy (taken by Luppi at al. to represent low-order dynamics) and their temporal synergy (high-order dynamics). By subtracting the redundancy rank from the synergy rank, Luppi et al. obtained a *synergy-redundancy rank gradient* of regions from the most low-order to the most high-order.

Applying this method to the mean information processing signature from each of the three diagnostic groups, we found this relative hierarchy (i.e. the relative positions of regions against each other) is essentially preserved in progression to AD, as can be seen in figure 15.

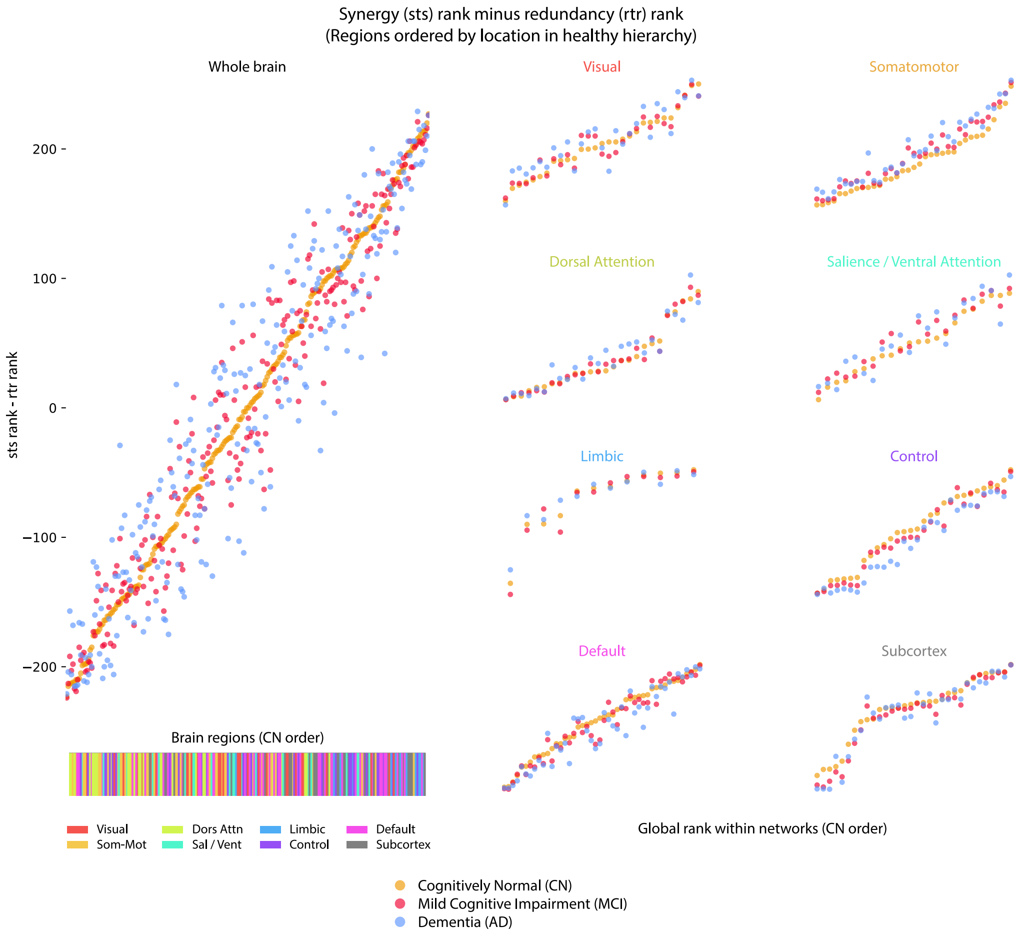


**Figure 15. Increased noise in the information processing hierarchy in AD.** Depicted is the synergy-redundancy rank gradient as constructed by Luppi et al. applied to the average information signatures across the CN group (orange), MCI group (red), and AD group (blue). Regions are ordered by their location in the CN hierarchy of information processing.

Correlating across regions to find a correlation between synergy-redundancy rank gradients of each group, we found very large correlations between all group hierarchies, with a slightly weakened correlation between the CN and AD groups. Confidence intervals were calculated using bootstrap resampling with $n = 1,000$ resamples.

| Groups | $\rho$ | 95% CI |
|---|---|---|
| CN with MCI | 0.975 | 0.951 – 0.979 |
| CN with AD | 0.937 | 0.894 – 0.948 |
| MCI with AD | 0.970 | 0.934 – 0.968 |

While correlation between the CN and AD hierarchy is slightly more degraded than between CN and MCI, these hierarchies are ultimately preserved, meaning that the broad arrangement of information specialisation is mostly unchanged (with respect to each other).

#### 6.11 Revalidation with Schaefer-116

All results were validated using an alternative extended Schaefer-116 atlas for additional verification. Much like the extended Schaefer 200 atlas, the extended Schaefer atlas used the original atlas of 100 regions due to Schaefer, with an additional 16 subcortical regions [61].

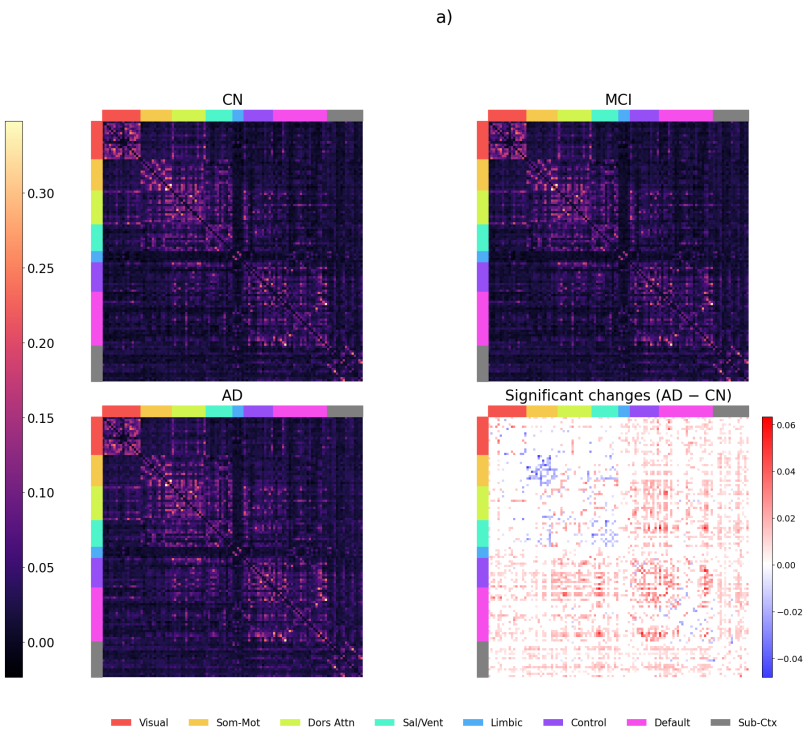

b)

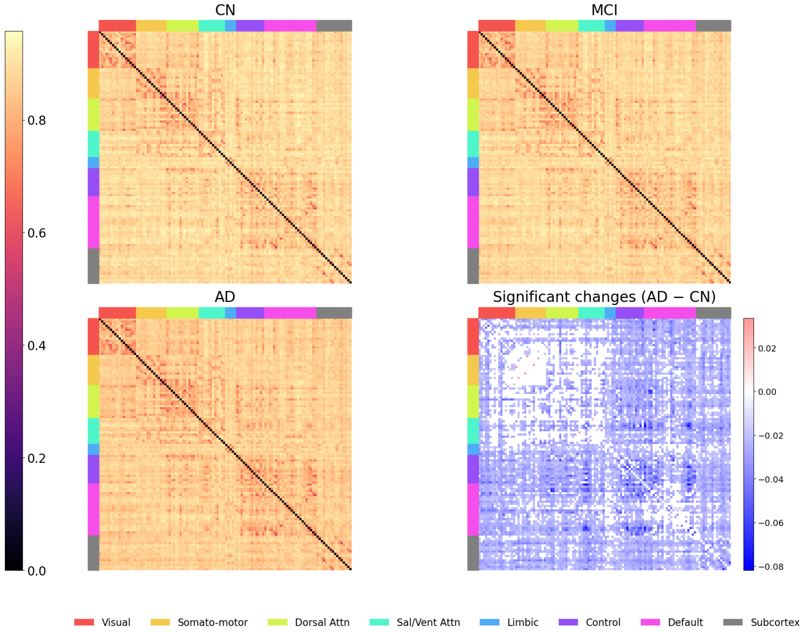


**Figure 16.** Matrices corresponding to the redundancy and synergy atoms from each coefficient tensor. As with the 232 matrix, regions are re-ordered into their 7 Yeo networks, plus additional subcortical regions. a) shows redundant information processing while b) shows synergistic information processing.

As with the extended Schaefer-232 atlas used in the main body of this work, somatomotor regions exhibited some decreased redundancy and minor changes to synergy. The rest of the pattern from Schaefer-232 is also replicated.

We also revalidated our intranetwork atom values. Significant changes are shown in figure 17. Changes to atoms on this atlas also replicate those seen in Schaefer-232.

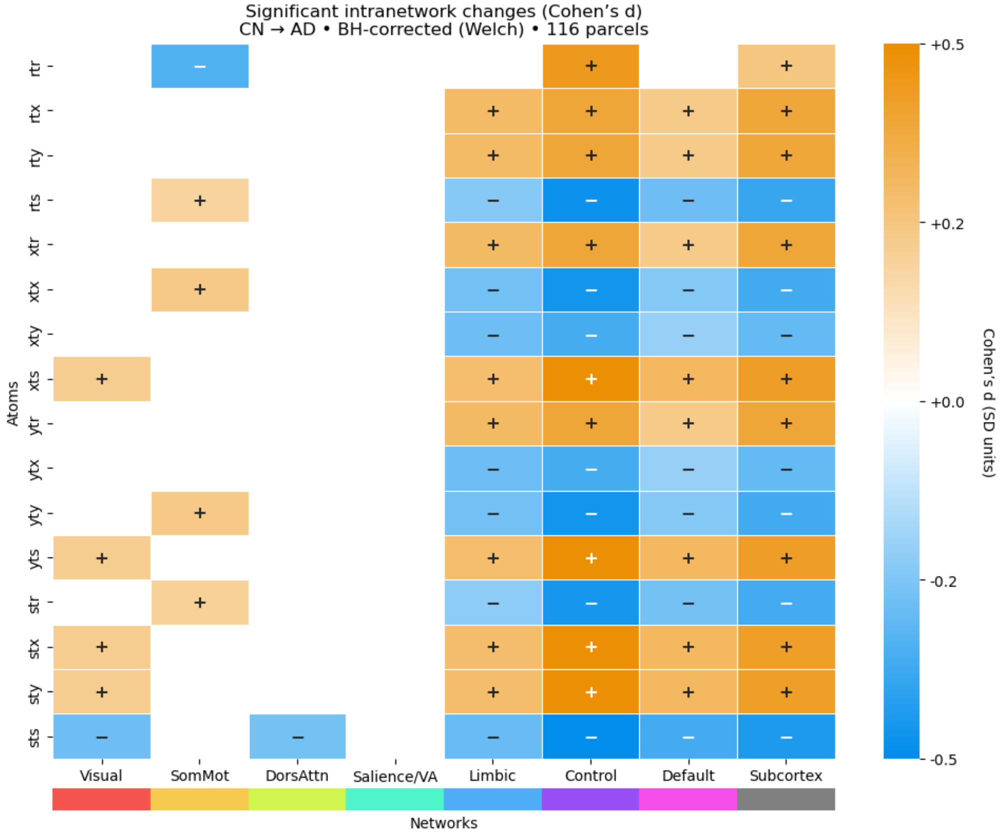


**Figure 17.** Atomic variation between CN and AD reported at the $\alpha = 0.05$ level with FDR correction, as per result 6.8, this time with 116 parcels.

#### 6.12 Coefficients found while regressing out confounds

As part of the analysis pipeline, we regressed out the effects of age, education, gender, and intracranial volume on each information atom, so as to more carefully assess the to underlying information processing profiles of each diagnostic group. The model also included an intercept. The values of the redundant and synergistic atoms are given in figure 18 for the intercept, the effect of being male, one year of education, one year in age, and each ml of intracranial volume. Due to computational constraints we did not threshold coefficients by testing the likelihood of being zero due to the complexity of an accurate statistical test. We report the coefficients here only for interest and illustration.

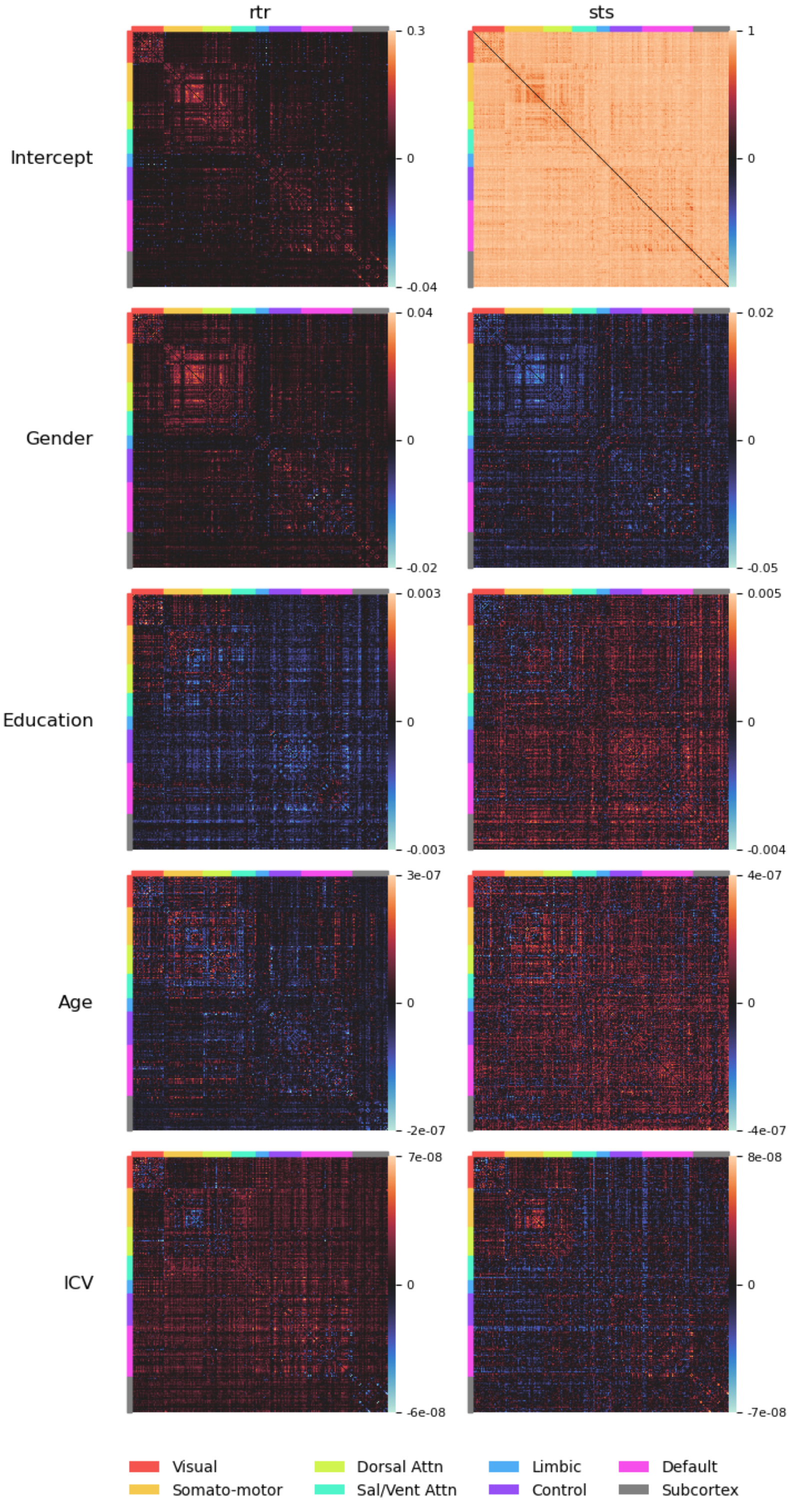
rtr
sts
Intercept
Gender
Education
Age
ICV
0.3
0
-0.04
1
0
0.04
-0.02
0.02
-0.05
0.003
-0.003
0.005
-0.004
3e-07
-2e-07
4e-07
-4e-07
7e-08
-6e-08
8e-08
-7e-08
Visual
Somato-motor
Dorsal Attn
Sal/Vent Attn
Limbic
Control
Default
Subcortex

**Figure 18.** Matrices corresponding to the redundancy and synergy atoms from each coefficient tensor. Regions are re-ordered into their 7 Yeo networks, plus additional subcortical regions. In all plots, black represents a contribution of zero, red a positive contribution, and blue a negative contribution. Education is measured in years, age in years, and total intracranial volume (ICV) in mL or $cm^3$. While included for reference, the effects of education, age, and ICV were mostly not significant at the $\alpha = 0.05$ level when accounting for the false discovery rate using a naive $t$-test. Gender as presented here corresponds to the effect of being male.

Results for the redundant and synergistic intercept reflect the patterns found in the original work by Luppi et al. [39], in that they appear to cluster according to network areas. Again we see intranetwork activity in the redundancy atom in the somatomotor network, with some other detail appearing in and between other combinations of networks.

The effect of being male appears to be mostly redundancy-bolstering, offering strongest increase in the visual, somatomotor and dorsal attention areas. When looking at the AD minus CN signature (c.f. figure 11), the effect of being male appears to be antagonistic to Alzheimer's decline.

Although not reaching the threshold for significance in our naive $t$-test, the same appears to be true of the education coefficient. Each year corresponded with an increase in redundancy in visual, somatomotor and dorsal attention regions. With 12 years in education, this effect becomes quite substantial, again potentially antagonistic to the change seen in AD against CN. Interestingly, age and intracranial volume had almost no effect on the information atoms, with units on the order of $5 \times 10^{-7}$ bits/year and $4 \times 10^{-8}$ bits/mL respectively. This might be due to the cohort already being observed in old age. Although ICV has been found to have some protective capacity in AD [27, 65, 66, 68], we did not find this reflected in our information measures here.

#### 6.13 Revalidation with CCS ΦID

In order to establish which results might be sensitive to the choice of ΦID method, we compared the MMI ΦID [2, 44] to another ΦID based on pointwise common change in surprisal due to Ince (CCS-ΦID) [29]. As with result 3.1, we applied $z$-scores to the MCI and AD groups using that region's redundancy and synergy in the CN group. In figure 19 it can be seen that CCS replicates the approximate $z$-scored pattern seen in result 3.1. In particular, both MCI and AD regions lost synergy when compared to the CN group. While almost all AD regions gained redundancy compared to the CN group, as with the MMI-ΦID, approximately half of regions lost redundancy in MCI compared to CN.

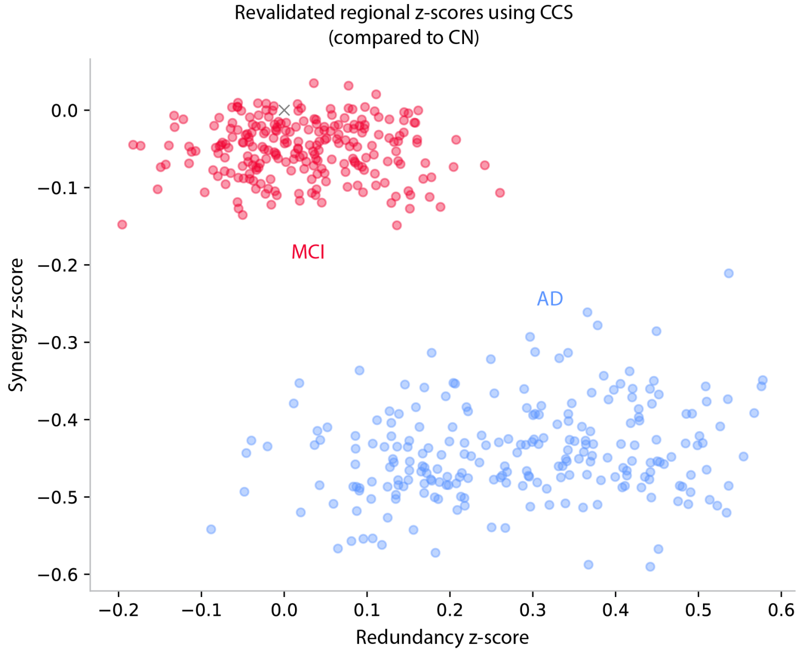


**Figure 19. CCS-ΦID exhibits similar synergy and redundancy changes in MCI and AD as MMI-ΦID.** For each region, the mean and standard deviation of synergy and redundancy. The corresponding regions for the MCI and AD groups had their synergy and redundancy $z$-scored against the CN mean and standard deviation. Red: $z$-scored regions from the average MCI signature of information representations. Blue: $z$-scored regions from the average AD signature of information representations. The CN mean for all regions is highlighted with a grey cross.

As in figure 1, we subsequently coloured regions based on their Yeo network, before projecting onto the first principal component of variation across regions. This can be seen in figure 20. Interestingly, there is less regional variation across the brain in how much synergy was lost in MCI and AD compared to CN, with the first principal component of variation in MCI being essentially flat, and with a slight positive slope in the AD group.

As with the previous result using MMI-ΦID, regions are approximately arranged by their network, with somatomotor regions lying at one extreme of the first principal component and ECN and DMN lying at the other— a structural pattern seen in the MMI-ΦID results for both diagnosis and MoCA score.

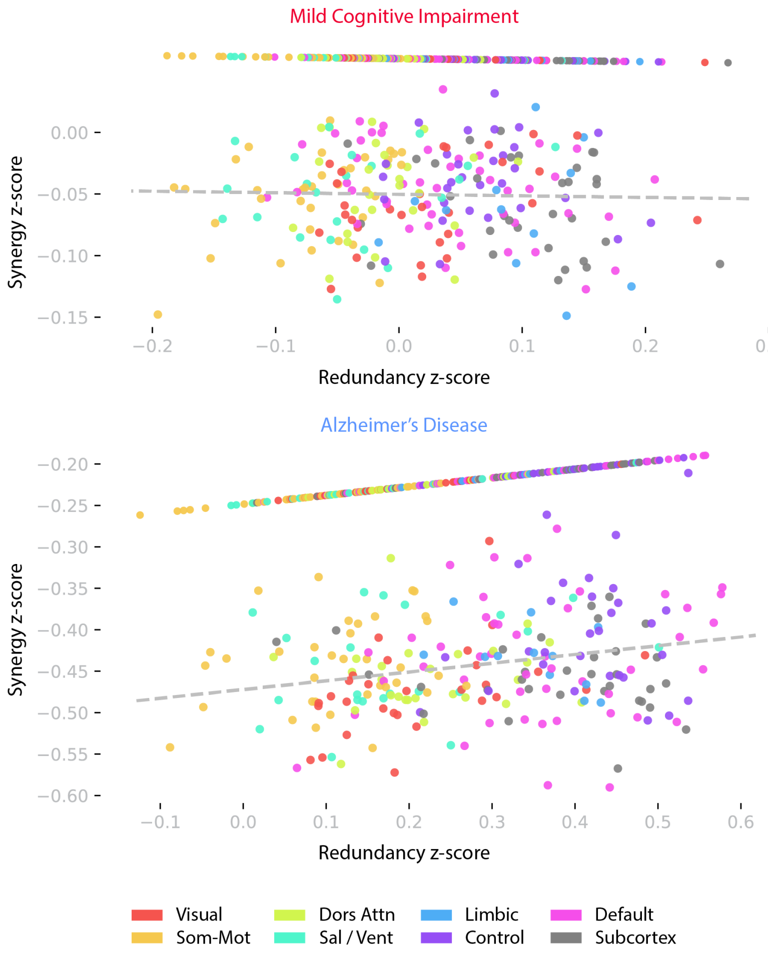


**Figure 20. CCS-ΦID shows similar network variation with less differentiated synergy profiles.** As in figure 19, $z$-scored regions from the average MCI (top) and AD (bottom) signatures are presented here, now coloured with reference to Yeo network. The first principal component of variation is also depicted, along with a projection of regions onto this axis.

In order to verify that the overall pattern of information representation seen was also similar in CCS-ΦID, we re-plotted matrices from 11, showing for each pair of regions $R_1$ and $R_2$, the amount of redundancy (figure 21) and synergy (figure 22) shared between them. We also tested, as with additional result 6.7, for significant differences between CN and AD pairwise processing, using an FDR-corrected $t$-test. In redundancy we found, as with 6.7, significant decreases in redundancy in the intranetwork somatomotor interactions (with some weaker decreases in salience regions), but increases in redundancy elsewhere.

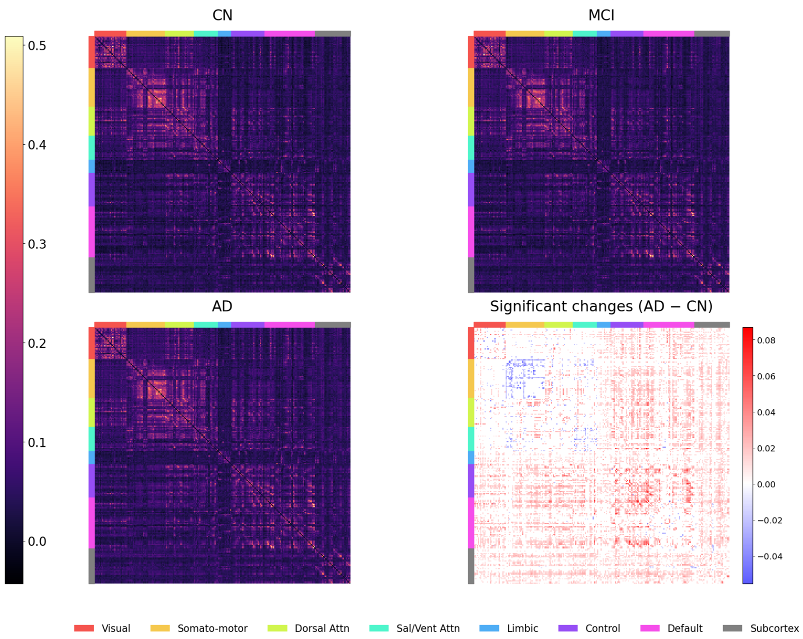


**Figure 21.** As with figure 11, for each pair of regions we plot the redundancy between those two regions across average matrices from the CN, MCI and dementia (AD) groups. Significant differences (found using an FDR-corrected $t$-test) between the CN and AD groups are highlighted in the bottom right, with significant increases shown in red and significant decreases shown in blue. Yeo networks are shown with different colours.

We repeated this with pairwise synergy, with results shown in figure 22. Of particular interest, we note that when using the CCS-ΦID, the significant decreases in synergy that were observed in MMI-ΦID are now absent. Overall $z$-scores in average synergy do appear to be smaller in magnitude in CCS-ΦID rather than in the MMI-ΦID, possibly accounting for this difference.

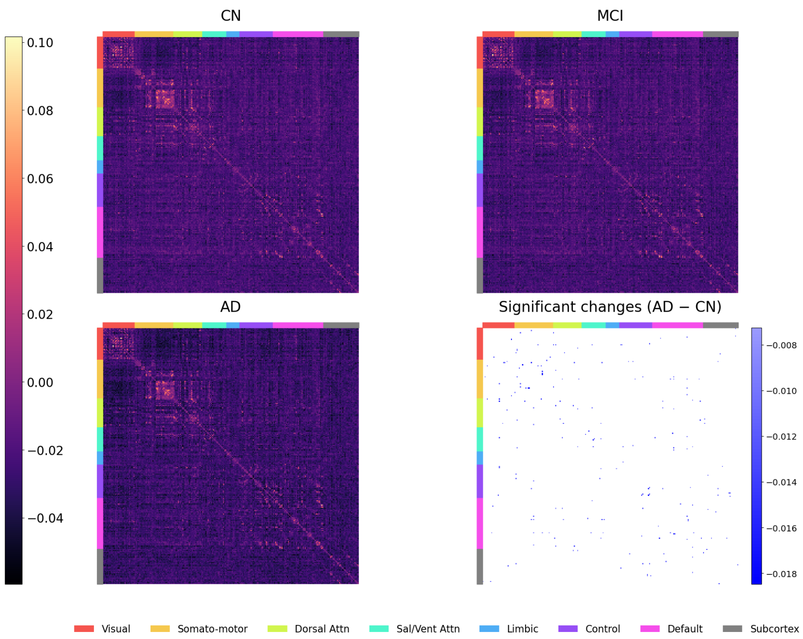

**Figure 22.** As with figure 21, but with CCS-ΦID synergy. For each pair of regions, the CCS-ΦID synergy between those two regions in plotted, with Yeo networks coloured accordingly. Significant decreases (although mostly absent) are coloured in blue, and were found by applying an FDR-corrected $t$-test.

Lastly, in order to verify that atomic changes were approximately the same as with MMI-ΦID, we looked at significant intranetwork differences as in additional result 6.8. Results are shown in figure 23. As with the MMI-ΦID, the somatomotor region shows oppositional behaviour to other networks when comparing the AD group to the CN group. As with the MMI-ΦID, the largest magnitude changes were observed in the ECN, with limbic, DMN, and subcortex similarly behind. It can be seen that these regions cluster approximately into the three groups: visual and DA regions, the somatomotor and salience regions, and the limbic, ECN, DMN and subcortical regions. Notably, the salience and DA networks showed more significant differences using CCS-ΦID compared to the MMI-ΦID.

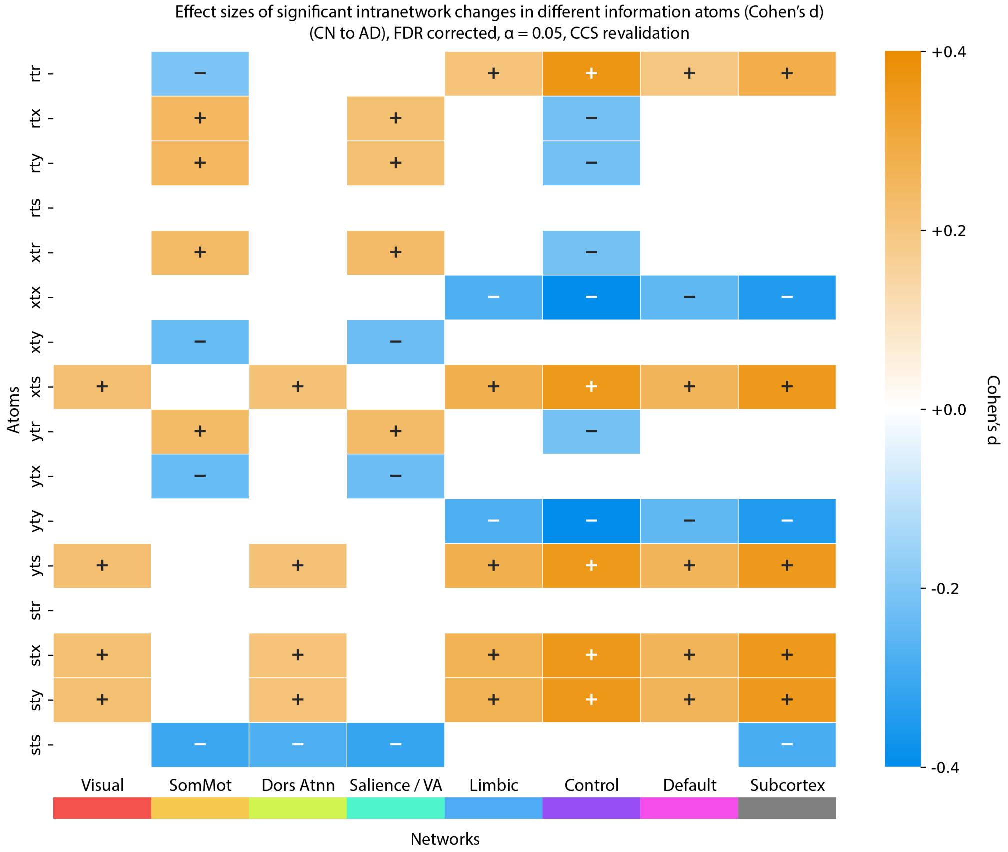


**Figure 23.** As like figure 12, we report atomic differences for intranetwork ΦID atoms. For each pair of regions within a single Yeo network (or in the subcortex), all CCS-ΦID atoms were calculated and averaged across that region only, giving an intranetwork measure of each atom. Significant changes between the CN and AD groups are reported at the $\alpha = 0.05$ level using FDR-corrected independent $t$-tests. For comparability we report the Cohen's $d$ effect size for each comparison, reporting significant increases in orange and significant decreases in blue.
